# RNA polymerase redistribution and increased gene dosage support growth in *E. coli* strains with a minimal number of ribosomal RNA operons

**DOI:** 10.1101/2022.02.10.479894

**Authors:** Jun Fan, Hafez El Sayyed, Oliver J. Pambos, Mathew Stracy, Jingwen Kyropoulos, Achillefs N. Kapanidis

**Affiliations:** Biological Physics Research Group, Clarendon Laboratory, Department of Physics, University of Oxford, Oxford OX1 3PU, United Kingdom; Institute of Fundamental and Frontier Sciences, University of Electronic Science and Technology of China, Chengdu, Sichuan, 611731, China; Department of Biochemistry, University of Oxford, Oxford OX1 3QU, United Kingdom; Kavli Institute for Nanoscience Discovery, New Biochemistry building, University of Oxford, South Parks Road, Oxford, OX1 3QU, United Kingdom

## Abstract

Bacterial transcription by RNA polymerase (RNAP) is spatially organised. RNAPs transcribing highly expressed genes locate in the nucleoid periphery, and form clusters in rich media, with several studies linking RNAP clustering and transcription of ribosomal RNA (*rrn*). However, the nature of RNAP clusters and their association with *rrn* transcription remains unclear. Here we address these questions by using single-molecule tracking to monitor the subcellular distribution of mobile and immobile RNAP in strains with a heavily reduced number of chromosomal *rrn* operons (*Δrrn* strains). Strikingly, we find that the fraction of chromosome-associated RNAP (which is mainly engaged in transcription) is robust to deleting 5 or 6 of the 7 chromosomal *rrn* operons. Spatial analysis in *Δrrn* strains showed substantial RNAP redistribution during moderate growth, with clustering increasing at the cell end-caps, where the remaining *rrn* operons reside. These results support a model where RNAPs in *Δrrn* strains relocate to copies of the remaining *rrn* operons. We also show that *Δrrn* strains experience increased *rrn* gene dosage in rich media, minimising growth defects due to *rrn* deletions. Our study further links RNAP clusters and *rrn* transcription, and offers insight on how bacteria maintain growth in the presence of only 1-2 *rrn* operons.

## INTRODUCTION

Transcription, a central process in gene expression, is spatially organised in many organisms; this organisation is thought to increase the efficiency for RNA synthesis (1) and help cells adapt to different growth environments, nutrients, and types of stress. In eukaryotes, synthesis of ribosomal RNA by RNA polymerase I occurs in the nucleolus, a nuclear compartment (2); further, eukaryotic mRNA transcription occurs in spatially enriched foci called “transcription factories” (3), which contain RNA polymerase II clusters (4) with lifetimes correlated to the levels of mRNA synthesis (5). Some viral transcription systems are also spatially organised; e.g., RNA polymerases of poliovirus form planar arrays/lattices with hundreds of molecules (6).

Transcription has also been shown to be spatially organised in bacteria, where early studies using conventional fluorescence microscopy in fixed cells showed that fluorescent derivatives of RNA polymerase (RNAP) in *E. coli* and *B. subtilis* (7) form bright, diffraction-limited foci in rich media, but not in minimal media; these prokaryotic transcription foci have been likened to transcription factories (3,8). Subsequent studies using photo-activated localization microscopy (PALM), a super-resolution imaging method, provided further insight into RNAP spatial organisation; using PALM microscopy on fixed cells at different growth conditions, it was shown that RNAPs form large clusters with ~70 and >100 molecules in rich media, and smaller clusters with ~35 molecules in minimal media (9). Single-molecule localisation studies in live *E. coli* cells showed that RNAPs tend to co-localize with the nucleoid lobes, while being nearly absent from the ribosome-rich cell endcaps (10,11). Further live-cell work combining PALM and single-molecule tracking was able to distinguish between mobile RNAPs (i.e., RNAPs exploring the nucleoid for promoters), and immobile RNAPs, with the latter fraction including transcriptionally active RNAPs that localized primarily at the nucleoid periphery; this study also provided the first observation of RNAP clustering in living bacteria (12).

Surprisingly, subsequent localisation-based work (13) suggested that RNAP clustering remained significant even when transcription was supressed, and only decreased substantially when all transcription was inhibited by rifampicin, leading to the proposal that the underlying nucleoid (rather than high transcription activity) controls the organisation of these RNAP clusters. Recently, it was suggested that RNAPs in bacteria form “biomolecular condensates” (14) via liquid–liquid phase separation (LLPS), a phenomenon seen in many organisms (15–17), including bacteria (18–21); the condensates were shown to contain high-density RNAP clusters in fast-growth conditions, and were mediated by protein–protein interactions, offering LLPS as an alternative mechanism that drives RNAP clustering (22).

A central point of debate in the spatial organisation of transcription and the formation of transcription foci is the exact role of ribosomal RNA operons (*rrn*). Ribosomal RNA transcription (which involves 16S, 23S, and 5S rRNA) accounts for ~85% of all active transcription in fast-growing cells (23); such high transcription levels are essential for sustaining rapid synthesis of the ~55,000 ribosomes (10) needed per daughter per cell cycle during rapid growth (24). Notably, *rrn* transcription is much less prevalent in minimal media. In the genomic map of *E. coli*, most of the seven *rrn* operons locate near the origin of replication (*oriC*), and all *rrn* operons orient in the same direction as DNA replication (Fig. 1A); this chromosomal location leads to increased gene dosage for *rrn* genes.

**Fig. 1.**
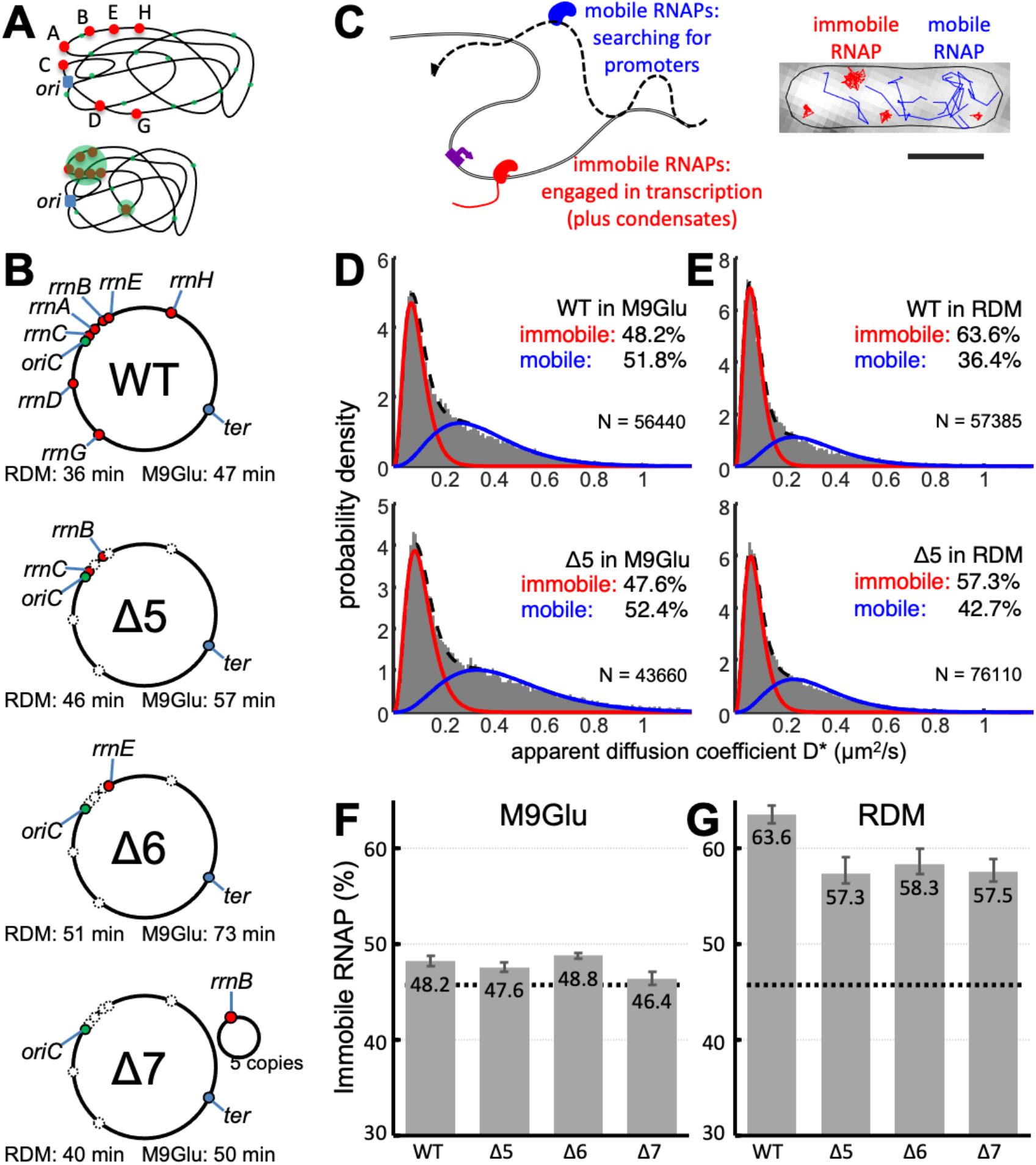
Measuring chromosome engagement of RNA polymerase at single-molecule level in *E. coli* strains with deletions of most chromosomal *rrn* operons. **A.** Models of RNAP distribution in cells based on observations of transcription foci and clustered RNAPs. Ribosomal operons are shown in red; RNAPs engaged in transcription are shown in green; origin of replication is shown in blue. Top: transcription-engaged RNAPs in minimal media are distributed throughout the nucleoid, with a bias towards the nucleoid periphery. Bottom: in rich media, transcription is concentrated on *rrn* operons, most of which appear in close proximity, forming large RNAP clusters. **B.** WT and *rrn* deletion strains used in this study; *rrn* loci are marked in the genomic map or in the supplemented plasmid. Doubling times of each strain in RDM and M9Glu media are also shown. **C.** Main mobility species of RNAP molecules and their characterisation using single-molecule tracking PALM. Scale bar, 1 μm. **D-E.** Histograms of apparent diffusion coefficient (D*) fitted with two-gamma distributions for WT and Δ5 in the M9Glu (**D**) and RDM media (**E**), along with the fractions of immobile and mobile RNAPs. N denotes the number of tracks per histogram. **F-G.** Fractions of immobile RNAP fractions for all strains in M9Glu (**F**) and RDM media (**G**). Dashed lines, median immobile RNAP fraction in M9Glu; error bars, S.E.M. from three individual measurements.

Transcription foci and RNAP clustering have been linked to *rrn* operons even during the first RNAP distribution studies (1,9), which raised the possibility that transcription foci involve multiple (perhaps all) *rrn* operons operating in close proximity and in a growth-dependent manner (since such foci were absent in minimal media), supporting a “bacterial nucleolus” model (2,24). Consistent with that model, Gaal *et al* measured the pairwise distance of *rrn* operons in single cells and found that 6 out of the 7 *rrn* operons in *E. coli* are in close proximity in 3D space (Fig. 1A, and Ref. (25)). Endesfelder *et al* also linked RNAP clustering to *rrn* operons, and suggested that clusters with 35-70 molecules represent single *rrn* operons, and large clusters (>100 molecules; 50-300 nm in diameter) represent super-clustered multiple *rrn* operons (9). These indirect links were supported by Weng *et al* (13), which directly showed that RNAP clusters were indeed colocalising with sites of high *rrn* transcription in rich media, while the formation of these clusters was independent of *rrn* transcription activity (13). The persistence of significant clustering despite the dramatic loss in *rrn* transcription was later attributed to LLPS (22).

Despite the progress in the understanding of spatial organisation of transcription, there are still many open questions. What is the link between RNAPs clusters and *rrn* operons, if any, in minimal media? To what extent LLPS contributes to RNAP clustering forming on *rrn* operons? What are the mechanisms that maintain the ability of cells to grow even when the number of chromosomal *rrn* operons is very small (1-2 copies)?

Here, we study the link between RNAP clusters and *rrn* operons by using single-molecule imaging and tracking (26,27) to obtain the RNAP spatial distribution and mobility in strains featuring deletions of most *rrn* operons (*Δrrn* strains; Fig. 1B). We show that, remarkably, in strains with only one or two chromosomal *rrn* copies, bacterial cells maintain the same level (~48%) of immobile RNAP (which mainly reflects RNAP engage in transcription) during moderate growth rates; immobile RNAPs in *Δrrn* strains move close to cell endcaps, suggesting that RNAPs relocate to the remaining *rrn* operons, which have a pole-proximal location. During fast growth in rich media, loss of most *rrn* operons decrease slightly the immobile RNAP fraction; for the strains with one and two remaining *rrn* operons on the chromosome, we show the presence of increased replication initiation in rich media, which will further increase the gene dosage for the remaining *rrn* genes. RNAPs retained its clustering in the *Δrrn* strains, whereas colocalization analysis showed a good correlation between RNAP clusters and *rrn* operons. In general, our results establish that loss of *rrn* operons is compensated through increased RNAP occupancy and increased gene dosage of the remaining chromosomal *rrn* operons. Our work expands our understanding of how RNAP is organised and allocated between transcription activities and how bacteria regulate their transcription to adapt to variations of their chromosomal content and growth environment.

## MATERIALS AND METHODS

### Bacterial Strains

The *rpoC:PAmCherry* WT strain carrying PAmCherry fused to the β’ subunit under the control of its native promoter used was built as described previously (9). The *Δrrn* strains were obtained by P1-transduction of the *rpoC:PAmCherry* gene in the *Δrrn* strains. The deletion strains were acquired from CGSC (*E. coli* Genetic Stock Center at Yale): SQ88 as Δ5rrn with *rrn*B and *rrn*C remaining, SQ110 as Δ6rrn with *rrn*E remaining and SQ2158 as Δ7rrn supplemented with plasmid-borne *rrn*B (pK4-16, based on pSC101, see also Ref (28)). For gene-dosage studies, *parS*^*pMT1*^ sites (29) were constructed by λ red recombination (30) around the *rrnC* or *rrnE* at the precise genomic positions used in (25). The parS-containing operons were introduced in *rpoC:PAmCherry* WT, Δ5, and Δ6 strains via P1 transduction. Foci visualization was obtained by constitutively expressing ParB-yGFP that selectively binds to *parS* sites from the pFH2973 (29); the strains used for this experiment were simply denoted by a parS suffix.

Cell growth rate measurements were performed using OD600 on a microplate reader (FLUOStar, BMG Labtech). Three separate measurements were carried out with individual blank media. The absorbance of OD600 was measured every 5 min for 16 hrs to generate the growth curves.

### Cell preparation for imaging

Strains were streaked onto LB plates supplemented with required antibiotics for each strain. For WT, we used 100 μg/ml ampicillin; for Δ5 and Δ6, we used 100 μg/ml ampicillin and 40 μg/ml spectinomycin, respectively; and for Δ7, we used 100 μg/ml ampicillin, 40 μg/ml spectinomycin and 50 μg/ml kanamycin. Single colonies were inoculated into LB and grown at 37 °C and 220 rpm for a pre-culture of 2 h, then diluted 1/250 into M9Glu media (1 X M9 media supplemented with CaCl2, MgSO4 and 0.2% glucose but *without* any additional vitamins or amino acids) or RDM (rich defined medium; Teknova) and grown at 37°C for overnight. Overnight cultures were diluted into fresh media and grown for >2 h at 37°C till early exponential phase (OD 0.1-0.2 for M9Glu culture, or OD 0.2 for RDM culture). 1.5 ml cell culture was centrifuged down, concentrated into 30 μl, and immobilized on 1% low-fluorescence agarose (BioRad) pads (supplemented with required M9Glu or RDM to keep media consistent). After immobilising the cells on agarose pads with fresh media, we monitored the RNAP localization using single-particle tracking on a PALM microscope and measured the apparent diffusion coefficient (D*) of RNAPs (see also Fig. 1C-D).

For fixed-cell colocalization experiments combining PALM with FISH, 1.5 ml of culture of the WT or ∆*rrn* strains carrying the rpoC:PAmCherry were spun down and then resuspended into 1 ml of PBS. One ml of 4% PFA was mixed 1:1 with the bacterial culture and incubated for 40 min with mild shaking on a nutator mixer at RT. After 3 washes with PBS, we added 500 μl of absolute ethanol to permeabilize the cells, and washed twice with PBS. We then immobilized 20 μl of cells on chitosan (31) housed in a self-adhesive gasket. For the FISH studies, 5 μM of the pre-rRNA probes carrying the sequence [Atto488]TGCCCACACAGATTGTCTGATAAATTGTTAAA-GAGCAGTGCCGCTTCGCT (13) was incubated with the permeabilized cells and incubated for 5 min at room temperature, then washed 3x with PBS. Cell were then imaged as discussed below.

### Single-molecule imaging of living cells

A custom-built single-molecule tracking PALM microscope (12) was used for the imaging of single RNAP-PAmCherry molecules and the detection of diffraction-limited *rrn* foci. Cells mounted on 1% agarose pads were imaged under bright-field illumination to perform cell segmentation. Prior to PALM imaging, pre-activated PAmCherry molecules were photobleached under continuous 561-nm excitation. Sparse photoactivation of the remaining population of PAmCherry was performed by continuous exposure to low intensity 405-nm excitation, such that the dataset consists of well separated single molecules. Under simultaneous excitation with a 561-nm laser, these photoactivated molecules fluoresce until permanently photobleached. Imaging was performed at a frame rate of 15 ms/frame for at least 30,000 frames, until the entire pool of RNAP-PAmCherry molecules had been imaged.

### Imaging and counting specific *rrn* operons by counting *parB* foci in live cells

Using parS-containing operons in strains WT, Δ5, and Δ6 supplemented with a plasmid constitutively expressing ParB-yGFP (see *Bacterial Strains*), we obtained estimates of the copy number of specific *rrn* operons in the WT and deletion strains. Cells were grown in RDM until OD600 reaches 0.2, immobilized on agarose pads, and imaged using 473-nm laser excitation and 500-ms exposures. The distribution of the number of foci per cell for the different strains was obtained after segmentation and localisation of the bright fluorescence parB foci using microbe, a plug-in for ImageJ (32). The results were then exported and plotted as a frequency histogram of the number of foci per cell.

### Two-colour colocalization assay

For FISH imaging of *rrn* foci with fixed cells, a 488-nm laser was used for 20 frames at 500-ms exposures. Bright field and pre-bleaching with a 561-nm laser were performed as for live cells. Excitation of RNAP-PAmCherry molecules was performed using a 561-nm laser for 90,000 frames at 15 ms/frame to capture the entire pool of RNAP molecules.

### Image processing and data analysis

Live-cell data were processed following published procedures based on a custom-written MATLAB software (12) for localizing single RNAP-PAmCherry molecules and tracking within a region-of-interest to distinguish different species. Histograms of apparent diffusion coefficient were fitted as two-gamma distributions, with the value for the immobile species fixed at 0.09 μm^2^/s. Comparison of different strains in defined growth media was done by collecting data in triplicates. Fixed cell data were processed using rapidSTORM as previously (9,33) to localize single RNAP-PAmCherry molecules while removing any repeated localisations in flanking frames.

### Clustering analysis of RNAP localisations

Clustering analysis of RNAP molecules localized by rapidSTORM was done using a MATLAB implementation of the DBSCAN algorithm (see Refs (9,34) and also Yarpiz page: Mostapha Kalami Heris, DBSCAN Clustering in MATLAB (URL: https://yarpiz.com/255/ypml110-dbscan-clustering), Yarpiz, 2015). From a Monte-Carlo simulation of localisations of RNAPs in M9Glu media and based on previous measurements in fixed cells (9), we determined the appropriate parameters for reliable clustering to be ε = 20 nm and MinPts = 4.

Pair correlation of RNAP localisations in live cells was performed as described previously (12). Briefly, the pair correlation function, g(r) was computed for all pairwise distances between the first localisations of each trajectory. For each cell, an equal number of simulated molecules were then distributed uniformly throughout the same segmented cell volume to generate another g(r) function which was used to normalise the experimental result. This normalisation compensates for the finite and asymmetric geometry of the cell.

### Pair correlation of RNAP clusters with *rrn* foci

Analysis of *rrn* foci in fixed cells was performed by bandpass filtering, followed by free elliptical Gaussian fitting for localisation. The pair correlation g(r) function was computed for all RNAP-*rrn* distances within the cell. A uniform distribution of the same number of molecules was generated throughout the segmented nucleoid area and used to normalise the experimental data. According to (35), nucleoid area covers ~56% of the overall cell area in M9Glu media, therefore we shrink both the cell length and width to ~75% to estimate the nucleoid area. A second uniform distribution was then generated and normalised using the same distribution as the experimental data to provide a visual guide for completely uncorrelated data, shown as the dashed line in the plots of the pair correlation function. Finally, the fraction of molecules found within 200 nm of *rrn* foci was calculated for both the experimental and uniform data.

### Heatmap plotting

Heatmaps of RNAP distributions were generated by pooling large populations of cells into two groups by cell length. Thresholds for cell length were selected such that the groups are composed of cells containing single and double nucleoids (Fig. S2). 2D histogram data of molecule location were obtained by binning data from normalised cell coordinates defined by the microbeTracker segmentation of each cell. Normalised data from all cells within the same cell length group were combined to produce heatmaps for mobile and immobile molecules. A difference map was generated by subtracting the mobile from the immobile distributions. Heatmaps were projected along their long and short axes to illustrate the RNAP distribution throughout the cell volume.

### Simulations

3D simulations of RNAP molecule locations were performed using Monte Carlo methods for the wild type, Δ5, and Δ6 strains. RNAP molecules were categorised into four populations; mobile, bound to *rrn* operons at the nucleoid periphery, bound in small clusters throughout the nucleoid, and ‘noise’ found throughout the cell volume. A total of 1800, 1800, and 1200 molecules were distributed between these categories in wild type, Δ5, and Δ6 respectively, consistent with experimental results in Fig. S9. A split of 48% immobile, 52% mobile molecules was simulated based on experimental data presented in Fig. 1.

A probability density function for non-*rrn* associated small RNAP clusters was obtained by fitting cluster size distributions obtained experimentally to an exponential function of the form y = ae^-bx^. The simulated population of non-*rrn* small cluster sizes are then generated by sampling from the inverse transform of this exponential probability density function, X = −(1/b)ln(1-(bu/a)), where u[0:1] is a set of uniformly distributed random numbers. The centres of non-*rrn* associated RNAP clusters were distributed uniformly throughout the nucleoid volume, around which RNAP molecules were distributed isotropically with a Gaussian radial density profile. Any molecules generated outside of the modelled cell volume were regenerated until a complete distribution was obtained.

The mobile population distributed uniformly within the nucleoid was generated via rejection sampling within a prolate spheroid volume positioned with a 150-nm separation between the nucleoid and cell poles. The same code was used to position the centres of small (non-*rrn*) clusters. A fraction of the mobile population was diverted to a population of localisation ‘noise’ that was generated uniformly across the entire cell volume via rejection sampling.

*rrn* operons were positioned along the pole-proximal periphery of each nucleoid. Simulations were performed with 7 operons/nucleoid in wild type, 2 in Δ5, and 1 in Δ6. The location of each cluster centre along the long axis which was weighted by the genomic distance of each *rrn* operon relative to *oriC*, defining a ring of possible locations around the nucleoid periphery. Candidate locations around this ring were proposed until a position was obtained with a minimum separation of at least 70 nm from other *rrn* operons. RNAP molecules were then distributed around each of these *rrn* cluster centres isotropically as described above for small non-*rrn* clusters. The proportion of immobile molecules associated with rrn operons was simulated across the range of 30 to 80%, with 60% to most closely matching the experimental data shown in Fig. 4. These 3D simulations were then projected into 2D, and analysed by computing the pair-correlation function g(r) for each cell. The process was then automated for 2000 cells to obtain the mean g(r) for the distribution.

### Estimation of the copy number of *rrn* operons and rRNAPs for a given growth rate

The total number of RNAPs engaged with the *rrn* operons (N_r_) for each strain was estimated by interpolation of the Nr values from Ref (36) [which were calculated using the expression N_r_ = r_r_/c_r_, where r_r_ and c_r_ are the overall rate of rRNA synthesis and the rRNA elongation speed (85 nt/s) as measured by Bremer and Dennis (37)] by fitting to a single exponential. The expected number of *rrn* operons in all strains for a given growth rate was obtained as in Bremer and Dennis (equation 9 in Table 5 of Ref (37)), by considering the growth rate of each strain, and the location of each *rrn* on map of the *E. coli* chromosome.

## RESULTS

### RNAPs remain heavily engaged with the chromosome despite deletion of most *rrn* operons

To clarify the relationship between RNAP clusters and *rrn* operons, we compared a well characterised *E. coli* strain carrying all 7 chromosomal *rrn* operons (“wild type”, WT) with strains carrying a drastically reduced number of *rrn* operons; these *rrn* deletion strains (*Δrrn*) were originally developed to study the link between *rrn* operon multiplicity and ribosome function (38,39).

Specifically, we studied a strain in which 5 out of 7 operons were deleted, leaving only *rrn*B and *rrn*C on the chromosome (Δ5, Fig. 1B); a strain in which 6 out of 7 operons were deleted, leaving only *rrn*E on the chromosome (Δ6, Fig. 1B); and a strain in which all 7 chromosomal *rrn* operons were deleted, and instead supplemented by a low copy-number plasmid (~5 copies per chromosome) containing a single *rrn*B operon (Δ7, Fig. 1B). To enable tracking of single RNAP molecules in cells, all strains contained a fully functional C-terminal fusion of the β’ subunit of RNAP with a photoactivatable mCherry (PAmCherry; in Refs (9,12); see also *Methods*).

To check the fitness of *Δrrn* strains relative to the WT, we monitored their growth in different media (Figs. 1B, S6); in general, the growth rates in the deletion strains correlated to the number of remaining copies of *rrn* operons, with Δ6 being the slowest growing. The WT strain displayed a 47-min doubling time at 37°C in minimal M9 media supplemented with 0.2% glucose (M9Glu; see *Methods*); in comparison, Δ7 grew marginally more slowly (50 min), Δ5 grew significantly more slowly (57 min), and Δ6 grew substantially more slowly (73 min). The growth rates of *Δrrn* strains in rich media followed a similar pattern; in rich defined medium (RDM), WT was the fastest growing (36 min), followed by Δ7, Δ5 and Δ6 (40, 46 and 51 min, respectively; for LB, see Fig. S6). In general, reduction in the number of *rrn* copies led to small-to-moderate decrease in the growth rate, presumably by affecting the rate at which different strains produce ribosomes (24,40).

To follow the RNAP mobility in live cells, we performed single-particle tracking of RNAP molecules using photoactivated localisation microscopy (PALM) on surface-immobilised cells, as described (Ref (12); see also *Methods*). The single-molecule tracks allowed us to calculate apparent diffusion coefficients (D*) for hundreds of RNAPs per cell (Fig. 1C-E; see also *Methods*) and construct D* histograms. The D* distribution for RNAP in WT cells grown in M9Glu was described well by two RNAP fractions with different mobility: a species (48.2% of all tracks) that remains immobile (D* of ~0.09 μm^2^/s), and a mobile species (51.8% of all tracks) with D* of 0.36 μm^2^/s (Fig. 1D, top; Fig. S1A, top). The immobile fraction includes RNAPs bound to the bacterial chromosome for several frames (12), which in turn correspond mainly to RNAPs bound to promoters and transcribed genes; the immobile species includes any RNAPs found in condensates, since they have been suggested to possess very low mobility (22). On the other hand, the mobile fraction corresponds to RNAPs interacting non-specifically and transiently with the entire chromosome during their promoter search (41).

To assess the effect of *rrn* operon loss on RNAP mobility in M9Glu, we compared the D* distribution of WT with those of *Δrrn* strains (Figs. 1D, 1F; Fig. S1C). In Δ5, and despite the deletion of 5 out of 7 chromosomal *rrn* operons, the D* distribution and the immobile RNAP fraction were essentially identical to those in WT (47.6%; Fig. 1D, bottom; Fig. 1F). The lack of any significant mobility difference compared to WT was also observed for both Δ6 and Δ7 strains (48.8% and 46.4%, respectively; Fig. 1F & Fig. S1B-C, left). These results establish that, despite the loss of most *rrn* operons, *Δrrn* strains show the same level of immobile RNAP as the WT.

To assess the effect of *rrn* operon loss on RNAP mobility in rich media (RDM), where the *rrn* operons should be much more heavily occupied by RNAPs than mRNA-coding genes (and where any RNAP condensates should be more visible, potentially accounting for ~30% of the immobile fraction; Ref. (22)), we performed similar RNAP mobility comparisons between WT and *Δrrn* strains in RDM. As we observed before (12), the immobile RNAP fraction in WT was ~63% (Fig. 1E, top; Fig. 1G), while Δ5 showed a ~6% decrease in the immobile fraction (57.3%; Fig. 1E, bottom; Fig. 1G, Fig. S1C, right); the results for Δ6 and Δ7 showed a similar decrease in immobile RNAPs (58.3% and 57.5%; Fig. 1G, Fig. S1B-C, right). In general, for all *Δrrn* strains grown in rich media, the immobile RNAP fraction stays surprisingly at the same high level (57-58%).

### DNA-bound RNAPs relocate to pole-proximal positions in *Δrrn* strains in M9Glu

To examine whether the deletion of most or all chromosomal *rrn* operons leads to any RNAP relocation within cells, and to gain insight regarding any redistribution between cellular RNAP pools, we examined the spatial distributions of mobile and immobile RNAPs in *Δrrn* strains via sorting single-molecule tracks using a D* threshold (Fig. 2A-B; see also Ref. (12) and *Methods*). To capture the average behavior for cells of similar size, we pooled the normalized positions of RNAPs from individual cells within different size ranges, and generated spatial heat-maps for both mobile and immobile fractions (see *Methods*).

**Fig. 2.**
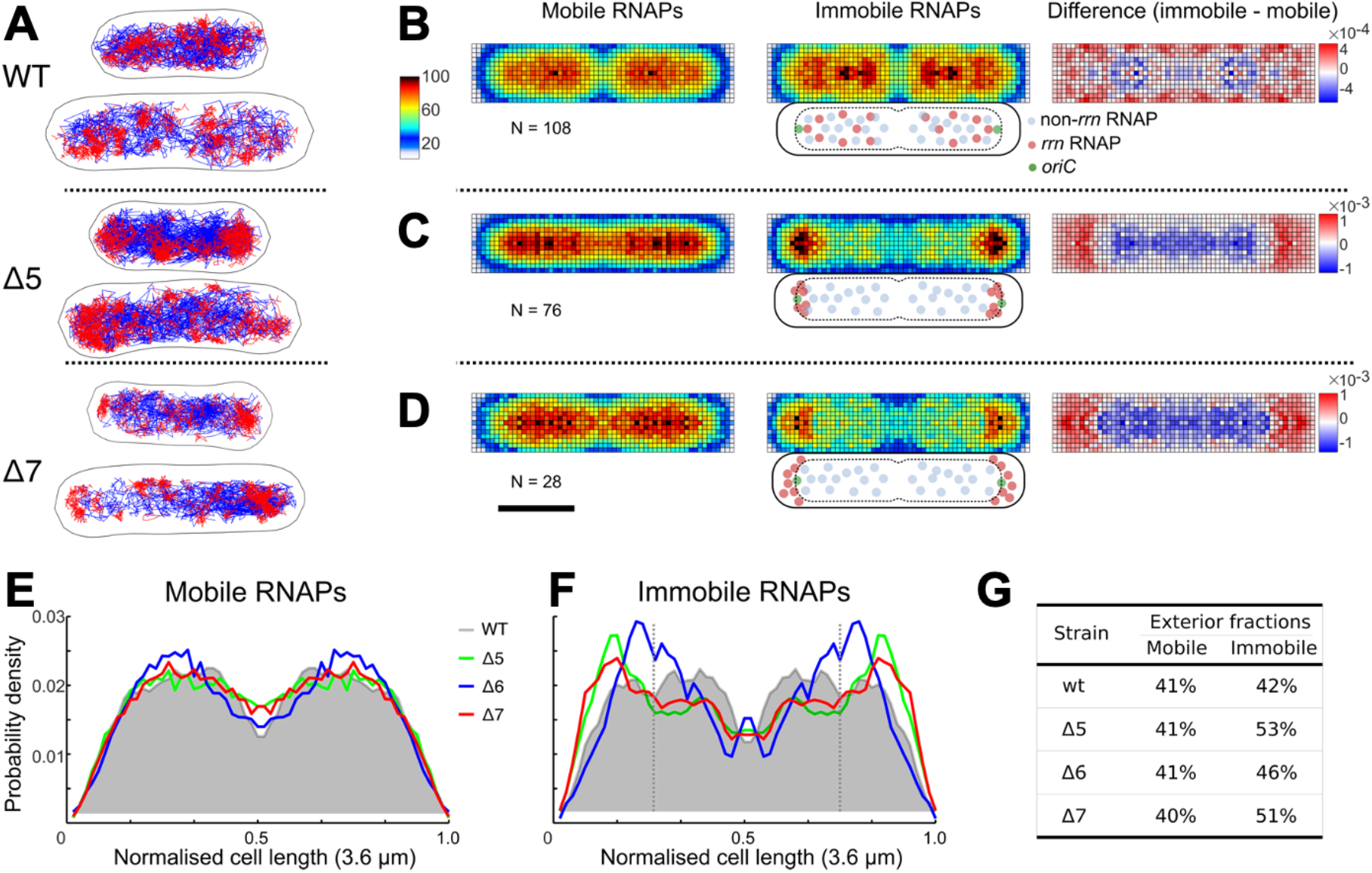
Immobile RNAPs in the Δrrn strains redistribute to polar-proximal regions in M9Glu media. **A.** Tracks of immobile (red) and mobile (blue) RNAP molecules for representative cell examples. The D* threshold for the track color-coding was 0.16 μm^2^/s. Scale bar (shown in panel D), 1 μm. **B-D.** Left and middle: heatmaps from multiple cells within 3.2 – 4.0 μm range of cell lengths for both mobile and immobile RNAPs. Right, difference map calculated by subtraction of mobile RNAPs from immobile RNAPs. Cartoons below the immobile RNAP heatmaps display the relative positions of *rrn*-related and non-*rrn*-related RNAPs relative to cell membrane, nucleoid and the *ori* region. Scale bar, 1 μm. **E-F.** Projections of mobile (**E**) and immobile (**F**) RNAP localisations along the long-axis of the heatmaps. The projection from WT is shaded grey. Dashed vertical lines indicate the 25% position along the long axis. **G.** The fractions of RNAPs localised in the exterior 25% region along the long axis (i.e., from 0 to 0.25 normalised cell length in Fig. 2F).

As we observed previously (12), the RNAP spatial distribution in WT cells with two nucleoids in M9Glu showed that mobile RNAPs localise throughout the nucleoid, essentially highlighting the nucleoid location (Fig. 2B, left), whereas immobile RNAPs tended to localize at the nucleoid periphery (Fig. 2B, middle); this redistribution towards the periphery for immobile molecules can be seen more clearly in the normalised difference heatmap between the two mobility fractions (Fig. 2B, right). Similar results were obtained for shorter cells, which carry only a single nucleoid (Fig. S2).

We then examined strain Δ5 to see how the deletion of 5 *rrn* operons affects the RNAP spatial distribution. The mobile RNAPs in Δ5 had a spatial distribution nearly identical to the WT one, i.e., spanning the entire nucleoid (Fig. 2C, left), reflecting the transient, non-specific interactions of this target-searching RNAP fraction with the nucleoid (41). In contrast, the spatial distribution of immobile RNAPs in Δ5 is substantially different to the WT one, with immobile RNAPs becoming much more concentrated at the pole-proximal edges of nucleoid (Fig. 2C, middle; see also the difference heatmap, Fig. 2C, right). For a clearer view of this RNAP relocation, we projected the heatmaps along cell length (Fig. 2E-F). While the projection for mobile RNAPs is similar, the projection of immobile RNAPs shifts from a fairly flat distribution centred at ~30% of cell length for WT, to a distribution with a peak at ~15% of cell length and Δ5 (green line, Fig. 2F); this shift is also reflected by the large increase in the RNAP fraction localised in the exterior 25% region along the long axis (53% in Δ5 vs. 42% in wild type; Fig. 2G); this increase corresponds to the relocation of ~10% of all immobile RNAPs.

A profile similar to Δ5 was observed for Δ7, which features only plasmid-borne *rrn* operons: the mobile RNAPs cover the entire nucleoid (Fig. 2D, left), while ~10% of the immobile RNAPs relocate to pole-proximal regions (Fig. 2D, middle and right; Fig. 2G), shifting the peak in the projection of immobile RNAPs to ~15% of cell length (Fig. 2F, red line). The Δ6 strain (with a single chromosomal *rrn* operon) also behaved similarly to Δ5 and Δ7 for both mobile and immobile RNAPs (Fig. 2E-G), with the main differences being the position of the new peak of immobile fraction, which appears at ~20% of cell length (Fig. 2E, blue line). Similar results were obtained for shorter cells (Fig. S3).

To explain our spatial distributions of immobile RNAPs, we need to consider that they contain several RNAP pools: RNAPs transcribing *rrn* operons (rRNAPs), RNAPs transcribing mRNAs (mRNAPs), and any condensate-associated RNAPs (cRNAPs). Removal of several *rrn* operons from the chromosome essentially releases many rRNAPs; since the immobile fraction for all 3 *Δrrn* strains does not change relative to WT, the released rRNAPs must join one or more of the 3 main pools of immobile RNAPs. Since the *Δrrn* strains do not have a substantial growth defect, it is likely that to maintain sufficient rRNA synthesis to support ribosome biogenesis, many (perhaps all) of the released rRNAPs are captured by the remaining *rrn* operons. Our results also show that a large fraction of immobile RNAPs in the *Δrrn* strains engage with the entire nucleoid (Fig. 2C-D middle); we attribute this fraction mainly to non-*rrn*-associated immobile RNAPs.

### Immobile RNAPs spread throughout the nucleoid in rich media

Since growth conditions influence dramatically the RNAP spatial distribution (1), we examined the spatial distribution in *Δrrn* strains growing exponentially in rich media (Fig. 3), a condition wherein cells need to accumulate high numbers of ribosomes (up to 70,000; see also Ref. (39)) and thus require high *rrn* expression (42,43). In rich media (RDM), WT cells divided every ~36 min, a growth rate that corresponds to ~2.7 chromosomes/cell on average, and a high copy of *rrn* genes (~23 *rrn*/cell; Refs (36,37)). The RNAP distribution in the WT strain in RDM showed that, as in M9Glu, mobile RNAPs explore the entire nucleoid, whereas many immobile RNAPs appear in clusters distributed throughout the nucleoid (see next section), with some enrichment at the nucleoid periphery (Fig. 3A; Fig. 3B; Fig. S4); this enrichment, especially visible along the short cell axis, is clear in the difference map between immobile and mobile populations, both for long and short cells (Fig. 3B, right; Fig. S4).

**Fig. 3.**
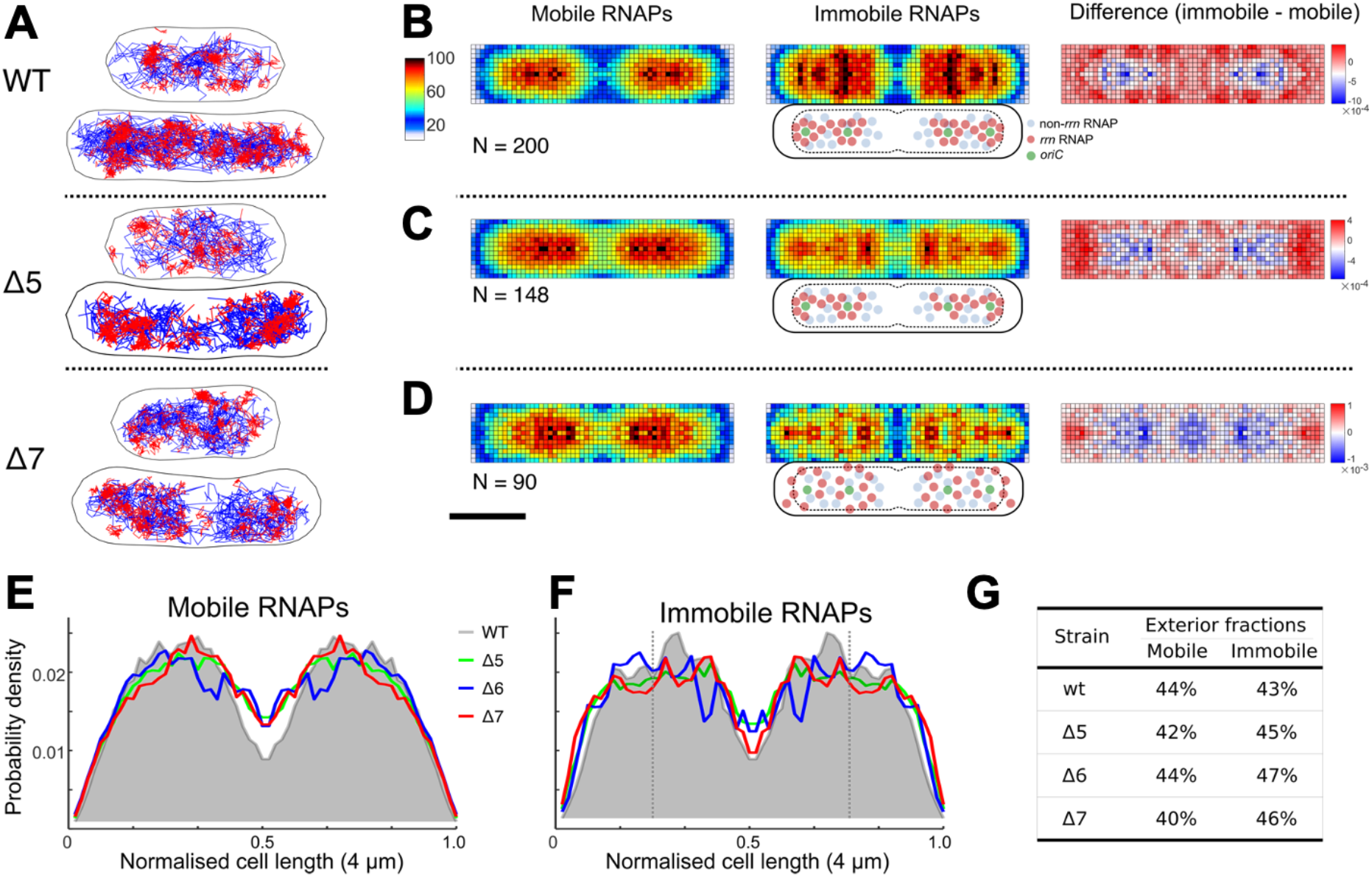
RNAP spatial distribution in the WT and *Δrrn* strains in rich media (RDM). **A.** Tracks of immobile (red) and mobile (blue) RNAP molecules for representative cell examples. The D* threshold for the track color-coding was 0.16 μm^2^/s. Scale bar (shown in panel D), 1 μm. **B-D.** Left and middle: heatmaps from multiple cells within 3.5-4.5 μm range of cell lengths for both mobile and immobile RNAPs. Right, difference map calculated by subtraction of mobile RNAPs from immobile RNAPs. Cartoons below the immobile RNAP heatmaps display the positions of *rrn*-related and non-*rrn*-related RNAPs relative to cell membrane, nucleoid and the *ori* region. Scale bar, 1 μm. **E-F.** Projections of mobile (**E**) and immobile (**F**) RNAP localisations along the long-axis of the heatmaps. The projection from WT is shaded grey. Dashed vertical lines indicate the 25% position along the long axis. **G.** The fractions of RNAPs localised in the exterior 25% region along the long axis (i.e., from 0 to 0.25 normalised cell length in Fig. 3F).

In contrast to its profile in M9Glu, the Δ5 strain shows a profile similar to WT for both mobile and immobile RNAPs, i.e., both populations are evenly distributed along the long axis of cells, and much of the immobile population appears clustered in single cells (Fig. 3A, 2^nd^ row). Intriguingly, and in contrast to the profile in M9Glu, no apparent relocation of immobile RNAPs to the poles is observed (Fig. 3C; Fig. S4), with the RNAP fraction in the pole-proximal region along the long axis being essentially identical for Δ5 and WT, both for mobile and immobile RNAPs (Fig. 3E-F). Broadly similar profiles and observations were seen in Δ7 (Fig. 3A, bottom; Fig. 3D) and Δ6 (Fig. S4; Fig. 3E-F), as well as in populations of short cells (Fig. S5). Notably, long cells feature a more polar and “nucleoid excluded” localisation of immobile RNAPs in Δ7 relative to the mobile population (Fig. 3D), likely reflecting the localisation of most plasmids.

To explain the spatial distributions of immobile RNAPs, we consider that, during fast growth conditions in rich media, cells contain multiple copies of the chromosome (9,44) and multiple sets of *rrn* operons, with the *ori*-proximal location of *rrn* genes further increasing the number of *rrn* copies (45); for example, for Δ5, we expect the group of long cells to have ~3 chromosomes and ~8 *rrn* copies on average (37). As in M9Glu, removal of several *rrn* operons from the chromosome releases many rRNAPs; since the bound fraction for all 3 *Δrrn* strains in RDM decreases only by ~6% relative to WT, the released rRNAPs must join one or more of the 3 main pools (rRNAPs, mRNAPs, and cRNAPs). We reason that, to maintain sufficient rRNA synthesis to support ribosome biogenesis (albeit at reduced growth rates), most released rRNAPs in *Δrrn* strains are re-captured by the remaining *rrn* operons (see also *Discussion*).

Our interpretation above is consistent with the location of RNAP clusters in single cells (e.g., Δ5 cells in Fig. 3A), which roughly map to the expected location of the 4 replication origins for cells of this size and growth rate (notably, the remaining *rrn* operons in Δ5 are proximal to *ori*). However, since the group of long cells covers a range of lengths (3.5-4.5 μm), and since the location of *rrn* operons varies for cells of different length, the average picture for the group of long cells is blurred and features fairly continuous distributions that do not reflect the localised nature of the clusters seen in single cells. On the other hand, Δ7 shows a clear profile of nucleoid exclusion for a large fraction of immobile RNAPs, suggesting that these represent RNAPs transcribing *rrn* genes on nucleoid-excluded plasmids. These *rrn*-centred RNAP pools are in addition to any pools of cRNAPs, although it is unclear whether cRNAP pools nucleate on the *rrn*-centred RNAP pools, or exist in isolation.

### RNAP clustering increases upon loss of most chromosomal *rrn* operons

The RNAP spatial distribution in *Δrrn* established that RNAPs in M9Glu relocate in pole-proximal regions, raising the possibility that relocation forms new clusters or enlarges smaller ones. To assess the level of RNAP clustering, we performed clustering analysis using the DBSCAN algorithm (Ref. (34) and *Methods*). Analysis of WT grown in M9Glu and subsequently fixed (Fig. 4A, top) showed that 29%, 11%, and 9% of clustered RNAPs were found in clusters with >35, >70, and >100 molecules, respectively (C>35, C>70, and C>100 species, Fig. 4B top) (9). For the Δ5 strain in M9Glu media, which shows an RNAP copy number per cell similar to that of WT (1767 vs 1844; Fig. S9), we detected a significant increase in all three species of large clusters (Fig. 4B), and notably for the C>70 population (17% for Δ5 vs 11% for WT), suggesting that RNAP redistribution increases the abundance of large clusters. Similar results were observed for Δ7, where despite a lower measured RNAP copy number for Δ7 (1285 vs 1844 for WT), the C>70 population increases to 20% (cf. 11% for WT; Fig. 4B, bottom).

**Fig. 4.**
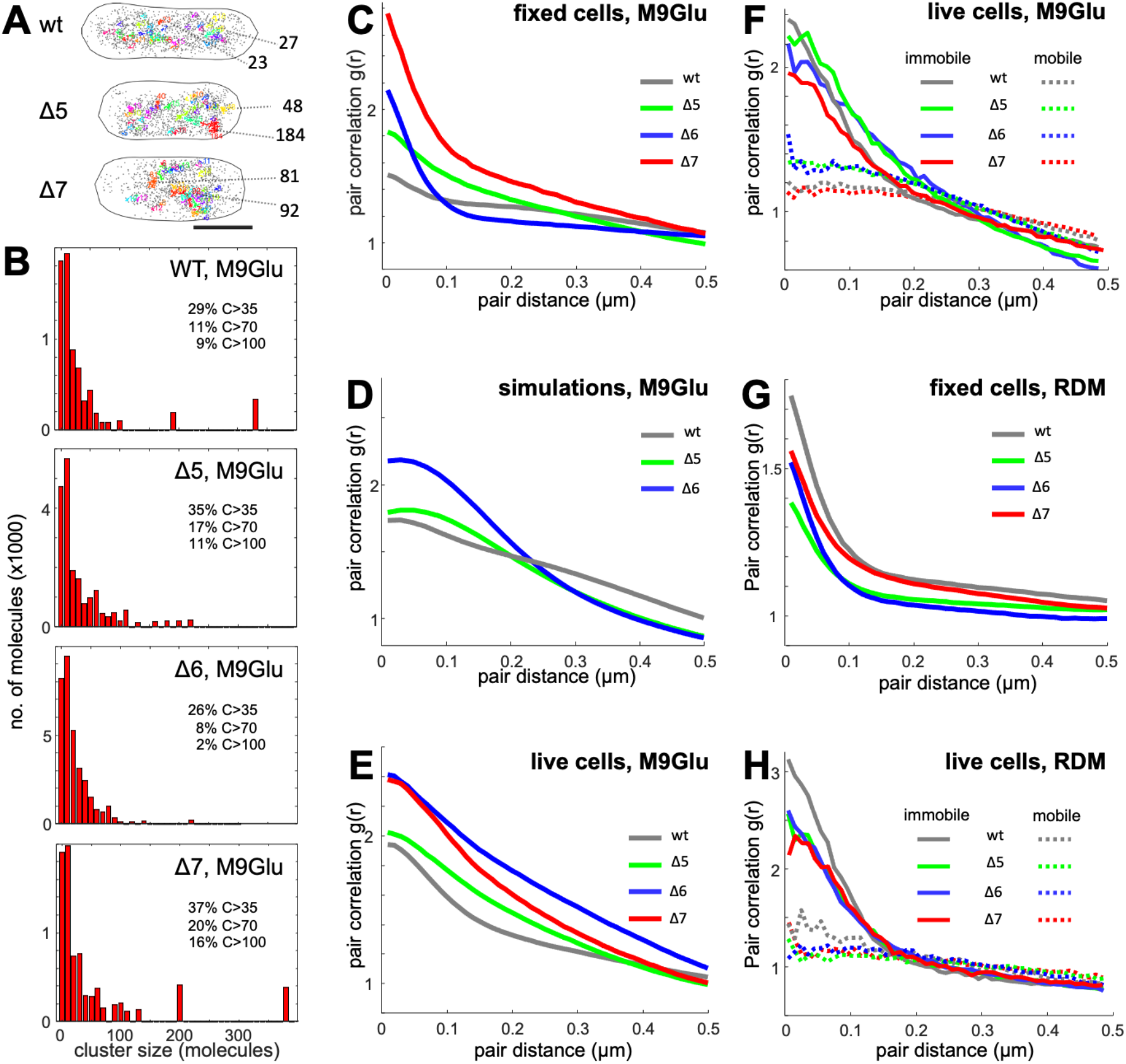
RNAP clustering and pair-correlation analysis of RNAP localisations in M9Glu and rich media. **A.** Representative examples of RNAP clusters in single fixed cells for WT and two *Δrrn* strains. RNAP clusters are displayed in different colors; non-clustered localizations are displayed as isolated grey points. Scale bar, 1 μm. Two clusters from each cell along with the cluster size (number of molecules) are shown as examples. **B.** Histograms of cluster size of RNAPs in fixed cells in M9Glu media. The fractions of large clusters of different size are listed alongside. **C-H.** Pair-correlation analysis of RNAP localisations in fixed and live cells of WT and *Δrrn* strains in M9Glu and RDM. **C.** Analysis in fixed cells of all strains in M9Glu. **D.** Analysis of simulated RNAP localisations for WT and two *Δrrn* strains in M9Glu. **E.** Analysis in live cells for all strains in M9Glu. **F.** Analysis of immobile and mobile RNAP tracks in live cells for all strains in M9Glu. **G.** Analysis of RNAP localisations in fixed cells for all strains in RDM. **H.** Analysis of immobile and mobile RNAP tracks in live cells for all strains in RDM.

Interestingly, there is a small reduction in large clusters in Δ6, in part due to a decrease in the copy number of RNAPs (1140; Fig. S9). These results are consistent with the deletion of most *rrn* increasing the degree of clustered RNAP due to relocation of RNAPs to the remaining *rrn* copies, all found in pole-proximal regions.

To gain another perspective to RNAP clustering, we performed pair-correlation analysis of the RNAP localisations (12), wherein the distances between all pairs of individual molecules are analysed and compared to a random distribution; a pair-correlation g(r) value of ≫1 for a range of inter-molecular distances indicates significant clustering, whereas g(r)~1 indicates a non-clustered distribution. Notably, the pair-correlation analysis is unaffected by differences in RNAP copy numbers per strain, and requires no optimisation in analysis parameters. We first examined RNAPs in fixed cells grown in M9Glu, and observed that RNAPs in WT show only slight clustering at distances within ~100 nm, whereas all three *Δrrn* strains showed much higher clustering within the ~100 nm range (Fig. 4C), with Δ7 being the most clustered strain. Pair-correlation analysis on simulated data for the two strains retaining chromosomal *rrn* copies (Δ5 and Δ6; see *Methods*) also showed that WT is expected to maintain the lowest level of RNAP clustering relative to Δ5 and Δ6 (Fig. 4D), consistent with our experimental results.

We also performed pair-correlation analysis in live cells in M9Glu. These experiments are complicated by any 3D motions of clustered RNAPs during the ~8 min of imaging; such motions will reduce the pair correlation and spread it out to longer length-scales; however, any persistent clustering should still be visible. We first analysed all RNAP localisations from the entire acquisition, and observed that all *Δrrn* strains show increased RNAP clustering relative to WT, with Δ7 and Δ6 being the most clustered (Fig. 4E); relative to fixed cells, the distance over which clustering is observed increases for all strains, as expected from the presence of some cluster mobility.

We then performed pair-correlation analysis on the mobile and immobile RNAP species in live cells in M9Glu; since we do not anticipate mobile RNAPs to be clustered (apart from exploring the entire nucleoid; as such, they do not fill the entire cell), this analysis should offer clearer views of the clustering of immobile RNAPs. Indeed, mobile RNAPs for all strains do not cluster (Fig. 4F, dotted lines); in contrast, the immobile RNAPs of both WT and *Δrrn* strains (Fig. 4F, solid lines) appear much more clustered than mobile RNAPs. Further, all strains show a similar level of clustering for immobile RNAPs. Taken together, our results indicate that in M9Glu, RNAPs become more clustered, consistent with the remaining *rrn* operons in *Δrrn* strains accommodating relocated RNAPs to compensate the loss of many chromosomal *rrn* operons.

In rich media, RNAPs in fixed cells of WT are more clustered than in M9Glu, whereas, in contrast, RNAPs in all *Δrrn* strains show reduced clustering relative to their levels in M9Glu, and relative to the WT in RDM (Fig. 4G). Pair-correlation analysis on the immobile RNAP molecules in live cells show similar differences between WT and the *Δrrn* strains (Fig. 4H); further, the levels of clustering in all strains exceed significantly the clustering seen in M9Glu. This result reinforces our qualitative observations of clustering in the discussion of the spatial RNAP distribution in RDM; the absence of prominent peaks in the projection of immobile RNAP localisations in the *Δrrn* strains are not due to the presence of highly distributed immobile RNAPs, but rather due to the presence of RNAP clusters with variable positions along the long-cell axis (due to the lack of synchronisation of the cells, and, in turn, due to variable positions of the remaining *rrn* operons).

### The expected cellular *rrn* copy number in *Δrrn* strains can sustain the measured growth rates in M9Glu, but not in RDM

To evaluate whether the remaining *rrn* copies can accommodate the number of RNAPs needed to sustain the measured growth rate (which is proportional to both cellular ribosome content [hence to the rRNA cellular content] and peptide elongation rate (46)), we estimated the copy number of *rrn* operons (37) and of *rrn*-associated RNAPs (36) for the growth rates of our strains (Table 1), and used them to estimate the average fractional occupancy of each *rrn* operon (Table 1).

**Table 1.** Estimated occupancy of *rrn* operons in WT and *Δrrn* strains as a function of growth rate.

| Strain | Media | Doubling time (min) | N<sub>rRNAP</sub> per cell* | *rrn* copies per cell | N<sub>rRNAP</sub> per *rrn* | RNAP occupancy of each *rrn* ** |
| --- | --- | --- | --- | --- | --- | --- |
| WT | M9Glu | 47 | 397 | 17.5 | 23 | 32% |
| $\Delta 5$ | M9Glu | 57 | 238 | 4.8 | 50 | 69% |
| $\Delta 6$ | M9Glu | 73 | 122 | 2.0 | 61 | 85% |
| WT | RDM | 36 | 805 | 22.0 | 37 | 51% |
| $\Delta 5$ | RDM | 46 | 420 | 5.6 | 75 | 104% |
| $\Delta 6$ | RDM | 51 | 320 | 2.5 | 128 | 178% |

For the methods used to obtain the estimates, see *Methods*.
\*Number of RNAP molecules transcribing *rrn* operons in the different strains.
\*\*100% occupancy is set to 72 rRNAPs/*rrn*.

For the WT strain (doubling time T of ~47 min in M9Glu), we expect an average of ~2 genome equivalents and ~17.5 *rrn* operons per cell (43), and ~400 rRNAPs (38), yielding an average of ~23 RNAPs/*rrn*. Considering a maximum *rrn* occupancy of ~72 RNAPs (using the average *rrn* occupancy at the maximal *E. coli* growth rate of 24 min), the operons operate at only ~32% of their full capacity, hence being far from saturation and having significant spare capacity.

Regarding Δ5 (T~57 min), we expect to have ~240 RNAPs engaged in *rrn* transcription and ~5 *rrn* operons per cell, leading to an estimate of ~50 RNAPs per operon, and ~70% of max occupancy; even if we use our experimental result of ~3.2 *rrn* foci/cell (see last *Results* section), and assume conservatively that a focus contains only one *rrn* operon, we recover an upper bound that does not exceed the *rrn* operon capacity. These estimates strongly suggest that the remaining *rrn* genes in Δ5 can accommodate the number of RNAPs required for the observed growth rate. Similarly, even for Δ6, the doubling time of 73 min can be maintained by an ~85% occupancy of the remaining ~2 *rrnE* copies per cell. In essence, the *rrn* transcription requirements for the growth rates of the deletion strains in M9Glu can be fulfilled by relocating RNAPs to the remaining *rrn* operons. Regarding Δ7, which has a growth rate similar to WT, the presence of ~10 copies of the *rrn*-containing plasmid (pK4-16, based on pSC101, see also Ref. (28)) also provides enough *rrn* copies to sustain the required levels of rRNA.

A more complex picture emerges for the *Δrrn* strains in rich media. In RDM, the WT (T~36 min) has ~22 *rrn* operons per cell and ~800 rRNAPs, with an average of ~37 RNAPs/*rrn*, which corresponds to operating at ~50% full capacity. As in the M9Glu case, since Δ7 has many copies (~15) of the *rrn*B-containing plasmid, it can maintain high *rrn* transcription levels despite the loss of all chromosomal *rrn*; indeed, Δ7 shows the smallest fitness cost (increase in doubling time) amongst the *Δrrn* strains.

However, in the case of Δ5 (T~46 min, ~420 RNAPs), we expect ~5.6 *rrn* per cell based on the measured growth rate, hence ~75 RNAPs/*rrn*, which exceeds the maximum capacity by ~5%; this means that Δ5 is at the limit of being able to sustain growth by fully loading all remaining *rrn* operons with RNAPs. This limit is substantially exceeded in Δ6, the slowest growing strain in RDM (Τ~51 min), which requires ~320 rRNAPs to maintain its growth rate, corresponding to ~130 RNAPs/*rrn*, which is physically impossible.

This result strongly suggests that mechanisms other than simple RNAP relocation to the number of *rrn* copies expected purely on the basis of growth rate are needed to explain the ability of Δ6 (and, possibly, of Δ5) to sustain the observed growth rate in RDM.

### *rrn* transcription in rich media for *Δrrn* is further compensated by increased gene dosage

Our estimates of RNAP occupancy of *rrn* in RDM clearly showed that, at least for the Δ6 strain, the cells cannot sustain the measured growth rate purely on the basis of the cellular number of *rrn* operons expected for the measured growth rate. This led us to hypothesise that more *rrn* are generated, likely due to increased replication initiation frequency. To explore this hypothesis, we used imaging to measure the number of *rrnE* copies in WT and in Δ6 growning in RDM; the *rrnE* copies were visualised as fluorescent foci (due to the binding of fluorescent ParB molecules) forming on *parS* sequences inserted in chromosomal positions adjacent to the *rrnE* operons in WT and Δ6 (see *Methods*).

Fluorescence imaging of the WT strain, followed by an automated analysis of foci (see *Methods*), showed that most cells have 1-4 bright foci (Fig. 5A), with the number increasing with cell length (Fig. S13E), and an average of ~2.8 foci per cell (SD~1.24; Fig. 5B), with the distribution width reflecting the fact that cells throughout the cell cycle are included in the analysis. The average number of foci is in very good agreement with the ~3.3 *rrnE* copies in RDM expected on the basis of growth rate (T~36 min).

**Fig. 5.**
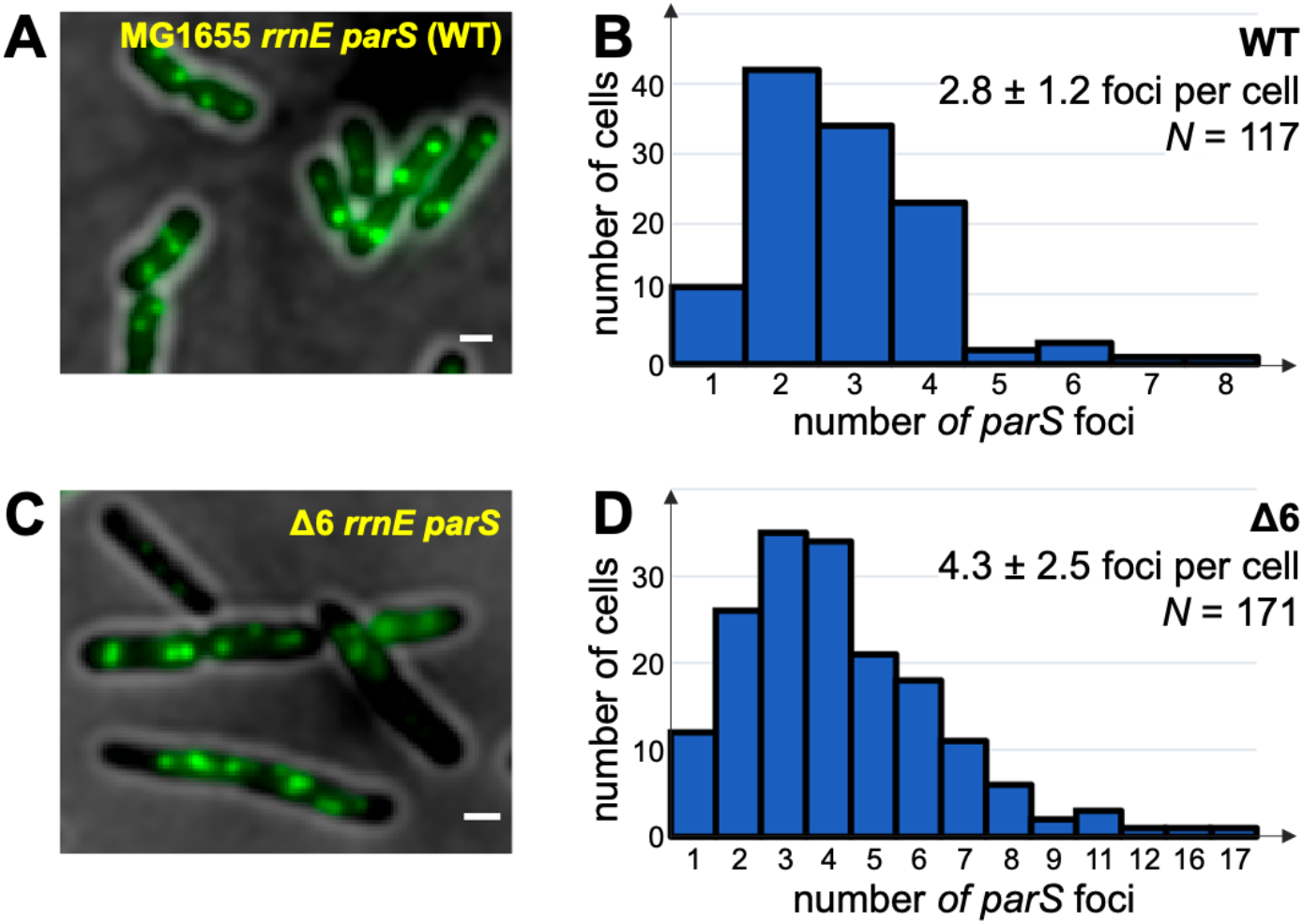
Imaging of *rrnE* operons shows increased gene dosage in RDM for Δ6 compared with WT. Fluorescence imaging of *rrnE* foci in the *rpoC:PAmCherry rrnE parS* strain harbouring plasmid pFH2973 expressing parB-yGFP. Cells were grown in RDM until OD600 ~0.2, immobilized on agarose pads, and imaged upon 473-nm laser excitation and using 500-ms exposures. **A.** Example of an image of cells containing parB foci at *rrnE* operons (appearing as bright green foci). Scale bar, 1 μm. **B.** Frequency distribution of the number of foci per cell for the WT strain, along with the mean and standard deviation of the distribution. Cells were segmented and foci were localised and counted using microbeJ (see *Methods*). *N* denotes number of cells. **C-D.** As in A and B, but for the Δ6 strain. The mean number of *rrnE* foci in Δ6 increases by ~70% compared to WT.

Similar analysis of the Δ6 strain showed that most cells carry a higher number of bright foci (up to 8 foci), for an average of ~4.3 foci per cell (SD~2.5; Fig. 5C-D). This increase leads to a number of *rrnE* copies that is ~70% higher than what is expected purely on the basis of growth rate (~2.5 copies), and leads to an estimated ~74 rRNAPs per operon, matching the maximum loading capacity of a *rrn* operon.

We performed a similar comparison between WT and Δ5 (Fig. S13A-D); considering the growth rates of the two strains, we were expecting ~28% more *rrnC* foci in WT compared to Δ5. However, the average of *rrnC* foci in WT is similar to that of Δ5 (4.8 foci in WT vs 4.9 in Δ5), consistent with the presence of additional replication initiation to increase gene dosage in the case of Δ5, albeit the effect is not as pronounced as in Δ6.

### Two-colour imaging reveals RNAP clusters correspond to *rrn* foci

To provide direct evidence for the link between RNAP clusters and *rrn* operons in the *Δrrn* strains, we performed two-colour colocalization assays by combining fluorescence in situ hybridization (FISH) imaging of *rrn* foci with single-molecule RNAP localisation in Δ5. Similarly to published work (13), we used a fluorescent FISH probe that targeted the 5′ leader region of the 16S precursor rRNA (pre-rRNA), which is absent from mature rRNA and ribosomes (see *Methods*). Signal from our FISH probe in fixed cells in M9Glu allows us to capture transcribing *rrn* foci, visualized as bright diffraction-limited spots in pole-proximal regions (Fig. 6A). When used in conjunction with PALM data, the relative colocalization identified active *rrn* clusters in Δ5 and estimated their copy numbers in M9Glu and RDM media to be ~3.2 ± 0.1 and ~6.8 ± 0.2, respectively (Fig. 6D). To record the position of *rrn* foci, its centroid was determined by a Gaussian fitting and superimposed on RNAP localizations and RNAP clusters; we observed that most RNAP clusters (especially large clusters containing >50 localisations; C>50) locate within 200 nm from the *rrn* centroid (Fig. 6A), suggesting significant co-localization between RNAP clusters and *rrn* foci.

**Fig. 6.**
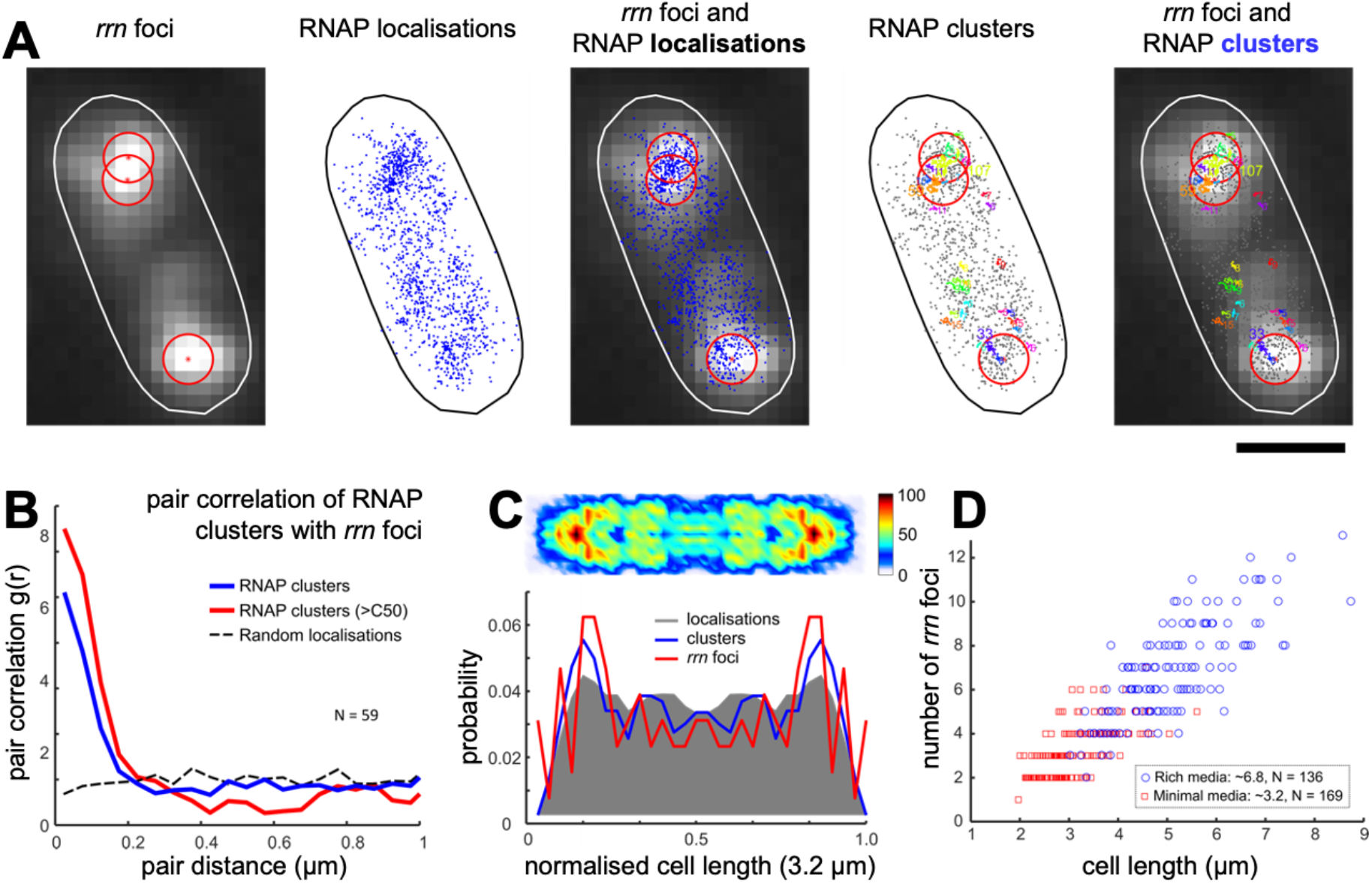
Co-localization of *rrn* foci with RNAP clusters in fixed cells. **A.** Representative example of a fixed cell showing FISH-labelled *rrn* foci in Δ5 along with RNAP localisations and clusters. Each centroid corresponds to one *rrn* focus obtained by Gaussian fitting and marked by a red asterisk and a 200 nm-radius red circle. RNAP localisations in fixed cells were analysed using DBSCAN to identify clusters; different clusters, along with their number of RNAPs, are shown in different colors. Scale bar, 1 μm. **B.** Pair-correlation analysis of *rrn* foci with all RNAP clusters, as well as large RNAP clusters (C>50) according to the pairwise distance show high co-localization between *rrn* foci with RNAP clusters and C>50 clusters: ~46% of RNAP clusters and ~77% of RNAP large clusters locate within a 200-nm radius of *rrn* foci (blue line), compared with ~23% obtained for simulated localisations randomly distributed throughout the nucleoid area (dashed black line). **C.** Heatmap of RNAP clusters and their projections (blue line). Projections of localisations and *rrn* foci are displayed in grey shaded area and red line, respectively. **D.** Scatter plot of *rrn* foci number in Δ5 in M9Glu and RDM as a function of cell length. Mean number of *rrn* foci is 3.2 ± 0.1 (SEM) in M9Glu, and 6.8 ± 0.2 (SEM) in RDM.

To quantify the degree of colocalization, we performed pair-correlation analysis between the *rrn* loci and RNAP clusters (either using all clusters with N>=4, or large C>50 clusters). Our results show a high correlation between *rrn* foci and RNAP clusters, supporting their colocalization. Specifically, we found that ~46% of all RNAP clusters, and ~77% of RNAP large clusters localize within 200 nm of *rrn* foci, compared with ~23% expected on the basis of simulated random RNAP localisations (Fig. 6B), which employs similar analysis algorithms as our pair-correlation analysis in Fig. 4.

To visualize the distribution of RNAP clusters, we generated the heatmap of RNAP clusters from the normalized positions of clustered RNAPs in Δ5 grown in Μ9Glu (Fig. 6C). Our results clearly show that RNAP clusters are concentrated at pole-proximal regions, a fact also reflected in projections along the cell-length axis. The projection of the normalized positions of *rrn* foci also display pole-proximal peaks, which highly overlap with the peaks of RNAP clusters (Fig. 6C). These results clearly establish the physical proximity of the RNAP clusters and *rrn* foci, and further support the suggestion that RNAPs relocate to the remaining *rrn* operons to sustain high levels of *rrn* transcription and largely maintain the growth rate achieved in the absence of any *rrn* deletions.

## DISCUSSION

The spatial organization of RNAP in bacteria has been a long-standing question ever since the first observations of transcription foci in cells grown in rich media (1–3,8,47–49), and the linkage between transcription foci and rRNA synthesis has remained controversial (1). Here, we applied super-resolution imaging and single-molecule tracking on strains with a heavily reduced number of *rrn* operons to elucidate the relation between RNAP spatial organization and rRNA synthesis, and study how cells redeploy their transcription machinery to sustain a healthy growth rate with only 1 or 2 chromosomal *rrn* operons. Notably, most bacterial species (~80%) have 1-4 *rrn* copies in their genome, with ~35% having just 1-2 copies (50); a large number of *rrn* copies enables the provision of high numbers of ribosomes per cell, which in turn allows bacteria harbouring a large *rrn* number to adapt faster to nutritional upshifts and switch to fast growth (and in general, respond more rapidly to changes in the nutrient availability) (50).

### RNAPs maintain their chromosome engagement in *Δrrn* strains by increasing the loading of the remaining *rrn* operons

Our RNAP mobility analysis showed that the fraction of immobile RNAPs, a proxy for the fraction of RNAPs engaged in transcription (plus any RNAPs involved in condensates), is surprisingly robust to the loss of 5 and 6 chromosomal *rrn* copies, as well as to the loss of all chromosomal *rrn* copies when cells are supplemented by a low-copy-number plasmid harbouring a single *rrn* operon. In M9Glu, a medium that supports a doubling time of 47 min in WT, about half of all RNAPs were immobile both for WT (as seen in Refs (12,51)) and all *Δrrn* strains. Even in rich media (RDM; supporting a doubling time of 36 min in WT), heavy loss of *rrn* operons led to only a modest decrease in the immobile RNAP fraction (63% for WT; 57-58% for the *Δrrn* strains).

The robustness of the RNAP immobile fraction to the loss of most chromosomal *rrn* genes raises the question of how RNAPs “released” from the deleted *rrn* operons redistribute to other immobile fractions, and how this redistribution minimises any growth-rate defects in *Δrrn* strains. Under our growth conditions in the WT strain (Fig. 7, top), we expect that *rrn* promoters are not saturated with RNAP; as a result, an increase in the concentration of available free RNA polymerase should lead to increased *rrn* promoter activities (46). Our results support a scenario where the remaining *rrn* copies in the cell are more heavily loaded by RNAPs (Fig. 7, middle).

**Fig. 7.**
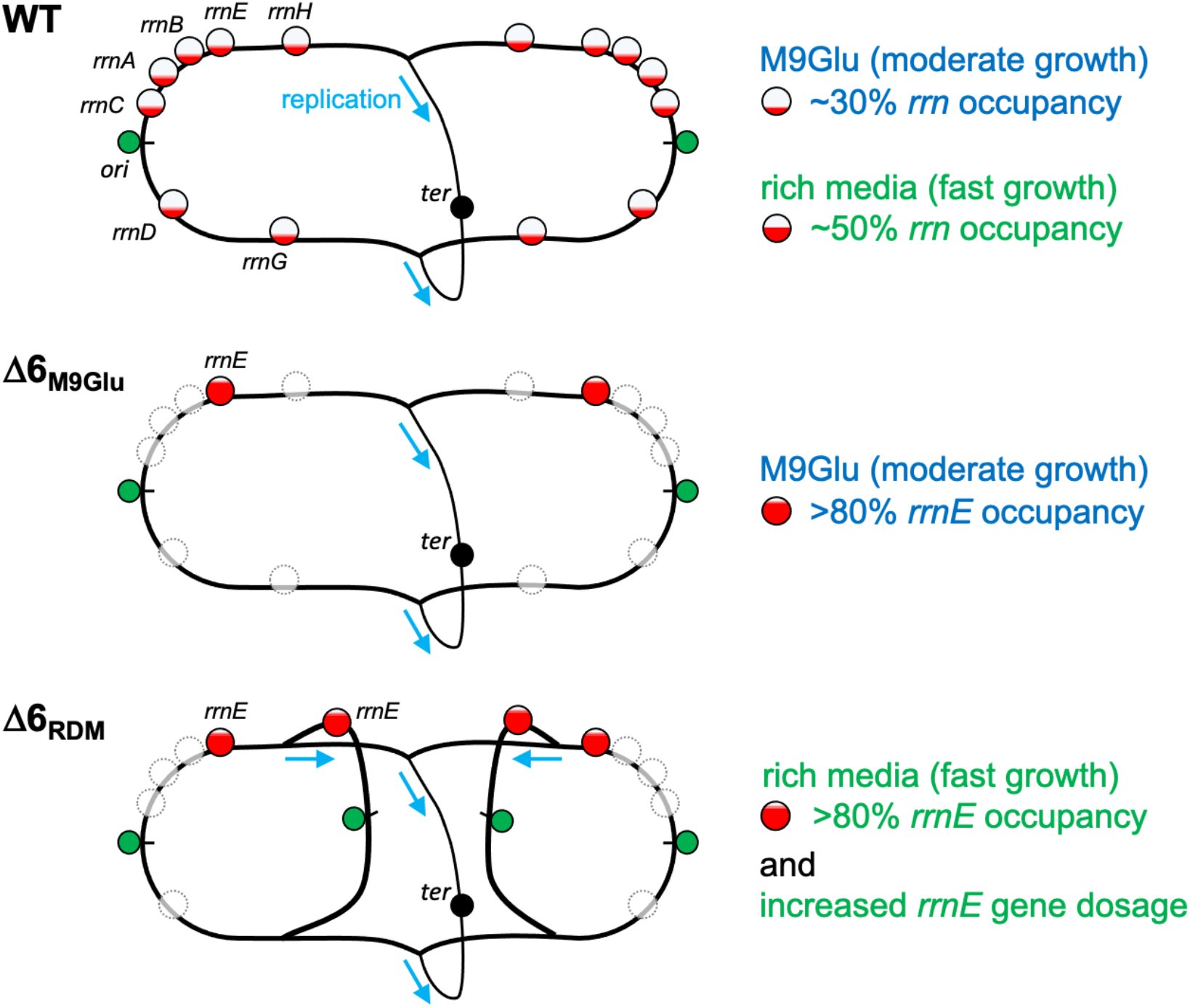
RNAP redistribution and increased gene dosage cooperate to maximise *rrn* transcription and minimize growth defects in *E. coli* strains with a small number of *rrn* operons. The *E. coli* strain with all 7 chromosomal *rrn* copies carries a large number of *rrn* copies (>15 copies) per cell either in M9Glu or rich media, with the operons being far from saturation (30-50% occupancy) in both media, and RNAP clusters forming on *rrn* spreading throughout the entire nucleoid. In contrast, in *rrn* deletion mutants lacking most *rrn* copies from the chromosome (e.g., Δ6), the remaining *rrn* copies are much more occupied by RNAPs (>80% in Δ6 in M9Glu) and RNAP clusters are more concentrated in the pole-proximal positions. More demanding growth conditions with regards to rRNA resources, such as in rich media, are facilitated by relative increase in replication initiation and a corresponding increase in the gene dosage for *rrn* (a ~70% increased dosage for Δ6), with RNAPs clusters spreading throughout the entire nucleoid.

Such RNAP re-distribution had been observed in a study of a more limited *rrn* deletion in rich medium LB; specifically, deletion of 4 *rrn* operons (Δ4) resulted in the remaining *rrn* operons accommodating ~71 RNAPs, an increase from 53 RNAPs/*rrn* in the strain with all operons intact, with the increased occupancy being linked to increased transcription initiation and elongation (38). Notably, the growth slow-down seen in Δ4 (~24 min for WT vs ~30 min for Δ4) is similar to that we see for Δ5 in RDM.

Our observations are consistent with the “saturation model” for the passive regulation on gene expression (46,52), and with studies showing that a 2- to 3-fold overexpression of the RNAP-σ70 holoenzyme leads to a 2-fold increase of *rrnB*P1 transcription, and a large increase in the tendency to transcribe *rrn* genes versus mRNA genes (53). In general, our results clearly show that the *rrn* promoters are competing very effectively against mRNA promoters.

### RNAP clusters do form on *rrn* operons

Our results clearly establish that RNAPs are more clustered in the *Δrrn* strains, and that RNAP clusters are in physical proximity to the *rrn* foci, adding further support to the proposal that RNAPs relocate to the remaining *rrn* operons to sustain high levels of *rrn* transcription and largely maintain the growth rate achieved in the absence of any *rrn* deletions. Our results are consistent with results from Weng *et al* (13), where it was shown that large RNAP clusters are maintained in a strain with a single *rrn* on the chromosome in rich media, as well as during low levels of transcription (13).

The presence of clustering on *rrn* (rRNAPs) does not exclude the presence of other forms of clustering, such as condensates (cRNAP) or heavy transcription on mRNA genes. It demonstrates, however, the ability of RNAP to redistribute and reprogram gene expression due to the cellular response to changes in the local environment or chromosome context (1,13).

### The location of remaining *rrn* operons dictates the location of the clusters in M9Glu

Considering the genomic map (Fig. 1B), the remaining *rrn* in the deletion mutants are either near *ori* (38), which is situated at pole-proximal regions along the cell long-axis (44), or on plasmids known to localise preferentially at the polar endcaps; these positions are consistent with the new peak of localisations (along the long cell axis) that appears in all 3 *Δrrn* strains (Fig. 2D). Relocation of released immobile RNAPs in MGlu in *Δrrn* to pole-proximal positions is thus dictated by the places of the remaining *rrn* operons.

The presence of a well-defined location for the relocated rRNAPs makes it unlikely that the released rRNAPs relocate to transcribe mRNA or join condensates not associated with transcription, since such RNAP pools are expected to have a much less localized position on the heat maps. In essence, it is the clustered rRNAPs that recruit cRNAPs (at least in the M9Glu case), and not the other way around.

### Heavy loss of *rrn* operons in rich media is further compensated via increased gene dosage

Our estimates of rRNAPs per *rrn* copy indicated that, to maintain a moderate growth rate in rich media, mechanisms additional to RNAP relocation are necessary; our measurements of fluorescent foci corresponding to specific *rrn* operons clearly demonstrated highly increased gene dosage, especially for the Δ6 strain. The increase in *rrn* copy numbers per cell is likely to result from increased frequency of replication initiation, thus linking gene expression with DNA replication (Fig. 7, bottom).

Such an intriguing connection has also been observed in *S. pneumoniae, E. coli* and other bacteria, in cases where stress induced by antibiotic treatments that target DNA replication (and stall replication forks) activates bacterial competence via a similar increased gene dosage of competence-related genes, which have an evolutionary conserved location near the replication origin (54).

In the case of the increased *rrn* gene dosage in Δ5 and Δ6 in rich media, we suggest that replication forks may be stalling due to co-directional collisions of the replication fork with RNAPs within *rrn* operons, since they will be fully occupied by elongating RNAPs and operate at maximum capacity; in contrast, replication initiation is unaffected, eventually increasing gene dosage for *ori*-proximal genes (Fig. 7, bottom). Such co-directional replication-transcription conflicts on highly transcribed *rrn* operons have indeed been reported (55), and depended on the presence of transcription and on rapid growth, during which >100 RNAPs per *rrn* operon were estimated to be present in *B. subtilis* (55).

## ACKNOWLEDGMENTS

The authors thank Dr. Olivier Espeli for providing the pFH2973 plasmid, the parSPMT1 genomic strain used for the *parS* strain construction and other molecular biology tools; CGSC stock center for providing the original *Δrrn* strains; the Oxford Synthetic Biology Doctoral Training Centre for access to a BMG Labtech microplate reader; and the MICRON Advanced Bioimaging Facility (supported by Wellcome Strategic Awards 091911/B/10/Z and 107457/Z/15/Z) for access to a PALM microscope in their facilities.

## AUTHOR CONTRIBUTIONS

A.N.K. and M.S. conceived the project. A.N.K., J.F, and H.e.S. designed the study. J.F., M.S., H.e.S., O.P., and J.K. performed experiments and analysed data. O.P. performed simulations, provided software, and analysed data. A.N.K. analysed data. J.F. and A.N.K. wrote the paper, and all authors had the chance to read and edit the paper.

## DATA AVAILABILITY

Movies and images of cells as well as localisation files for single molecules will be available upon request.

