## Supplemental Information for "RNA polymerase redistribution and increased gene dosage support growth in *E. coli* strains with a minimal number of ribosomal RNA operons"

### Supplemental Figures for Fan *et al* manuscript

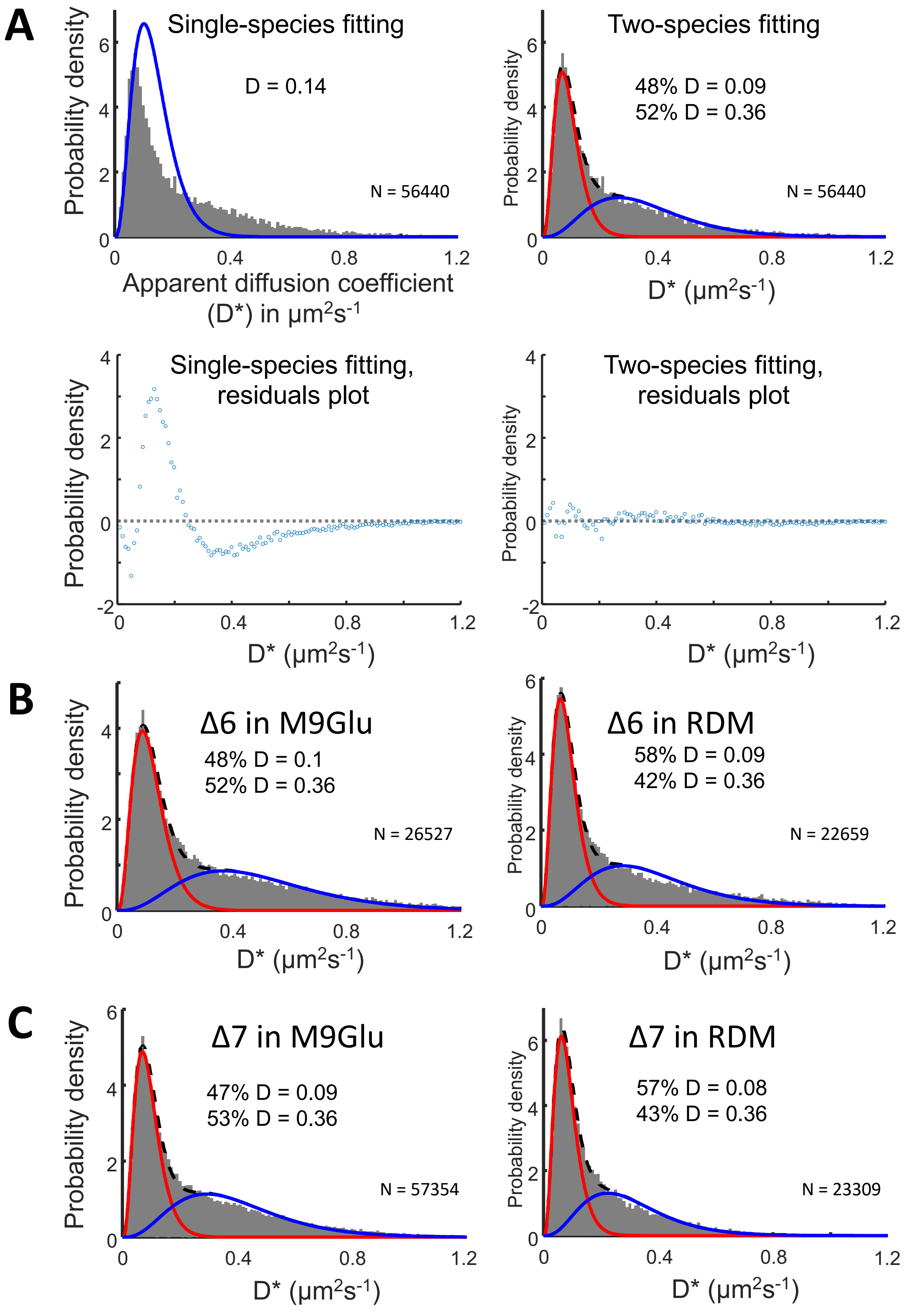


**Fig. S1. Distribution of diffusion coefficients of RNAP species in WT and deletion strains. A**. Top panels: fitting of the histogram of apparent diffusion coefficients (D*) with one or two diffusing species. Sample: WT strain in M9Glu media. Bottom panels: distribution of the fitting residuals. **B.** D* histograms fitted for Δ6 in M9Glu (left) and rich media (right) using two gamma distributions. The mean fractions of immobile (D* = 0.09-0.1 μm$^2$/s) and mobile RNAPs (D* = 0.36 μm$^2$/s) are listed. **C.** D* histograms for Δ7 in the M9Glu (left) and RDM media (right), fitted using two gamma distributions. Mean fractions are essentially as in panel B.

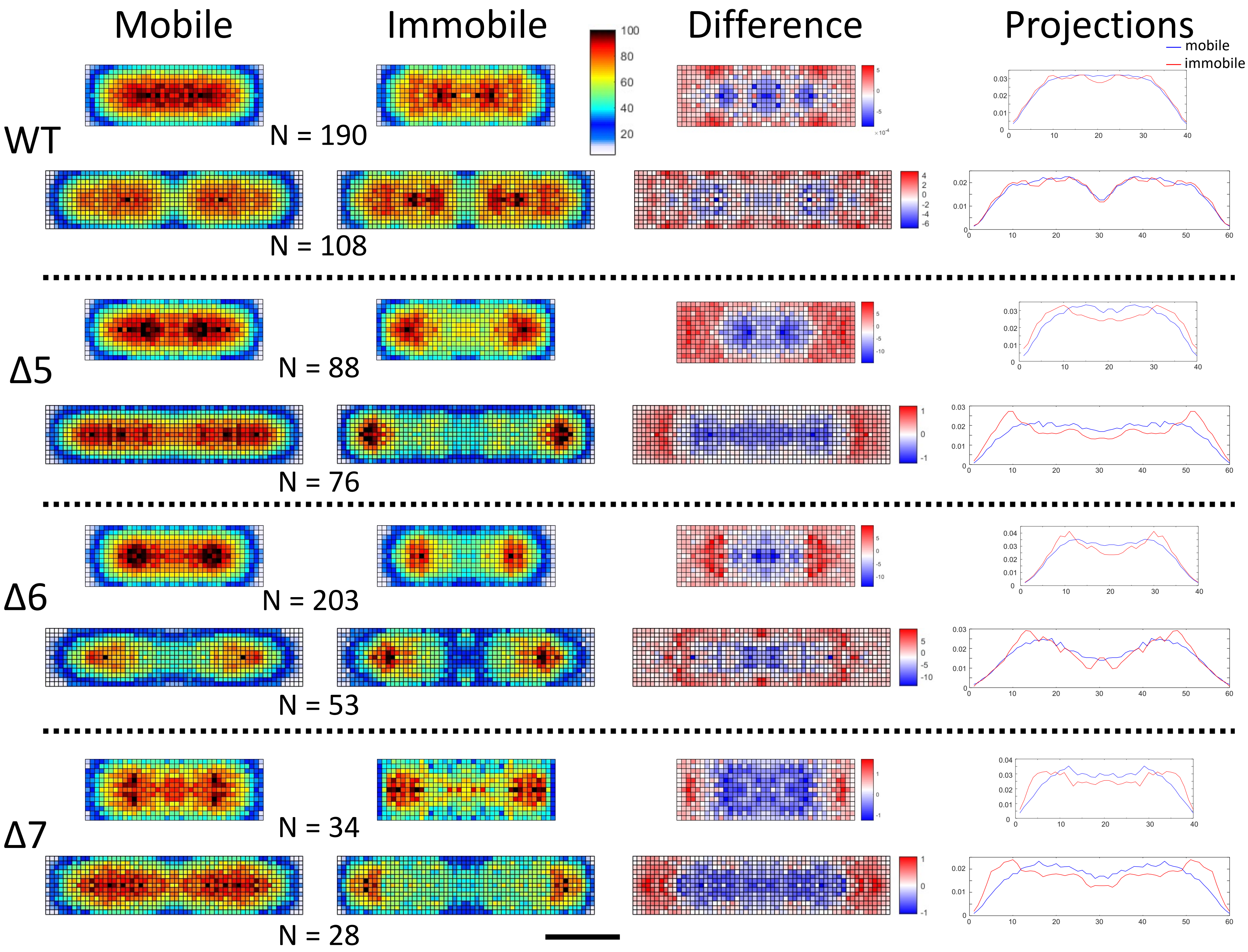


**Fig. S2. Heatmaps of mobile and immobile RNAPs and projections along the long-axis for all strains in M9Glu media.** For each strain, cells were sorted into a short group (top: 1.8-3 µm) and long group (bottom: 3.2-4 µm). In all *Δrrn* strains in M9Glu, immobile RNAPs redistribute to subcellular regions in close-proximity to the pole. Mobile RNAPs distribute throughout the nucleoid for all strains. The redistribution of immobile RNAPs is also reflected in the difference map (calculated by subtraction of mobile RNAPs from immobile RNAPs) and projection curves along the long axis of the cells. Scale bar, 1 µm.

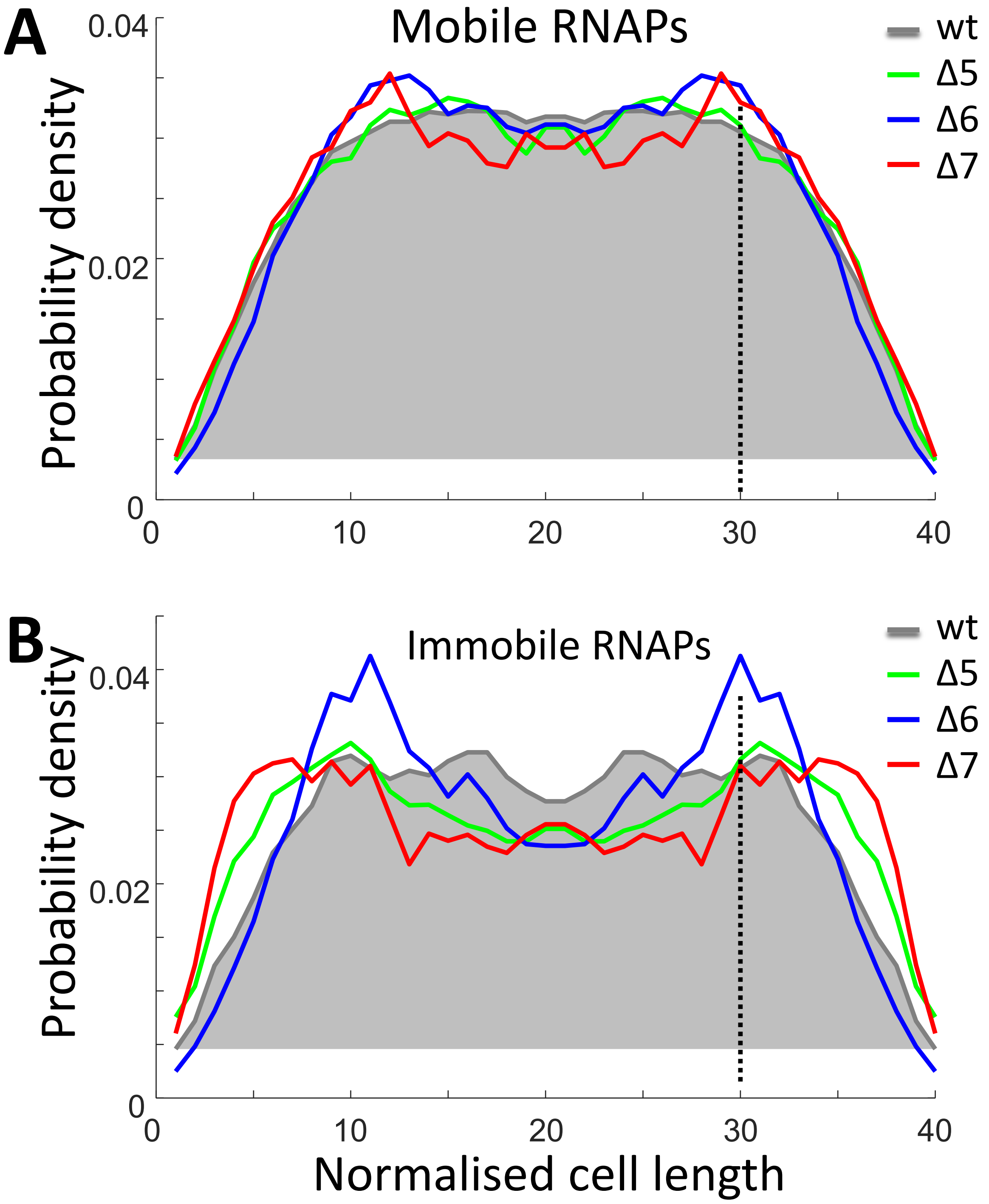


**C**

| Strains | Exterior fractions | |
|---|---|---|
| | Mobile | Immobile |
| wt | 37% | 39% |
| Δ5 | 37% | 47% |
| Δ6 | 34% | 40% |
| Δ7 | 39% | 50% |

**Fig. S3. Spatial distributions of the mobile and immobile RNAPs for short cells in M9Glu media. A-B**. Projections of mobile (**A**) and immobile (**B**) RNAPs along the long axis of the heatmaps for the short-cell group (1.8-3 µm length range) in M9Glu. WT projection is shown as a gray-shaded area. Dashed vertical lines indicate the 25% position along the long axis. **C**. Fractions of RNAPs localised in the exterior 25% region along the long axis.

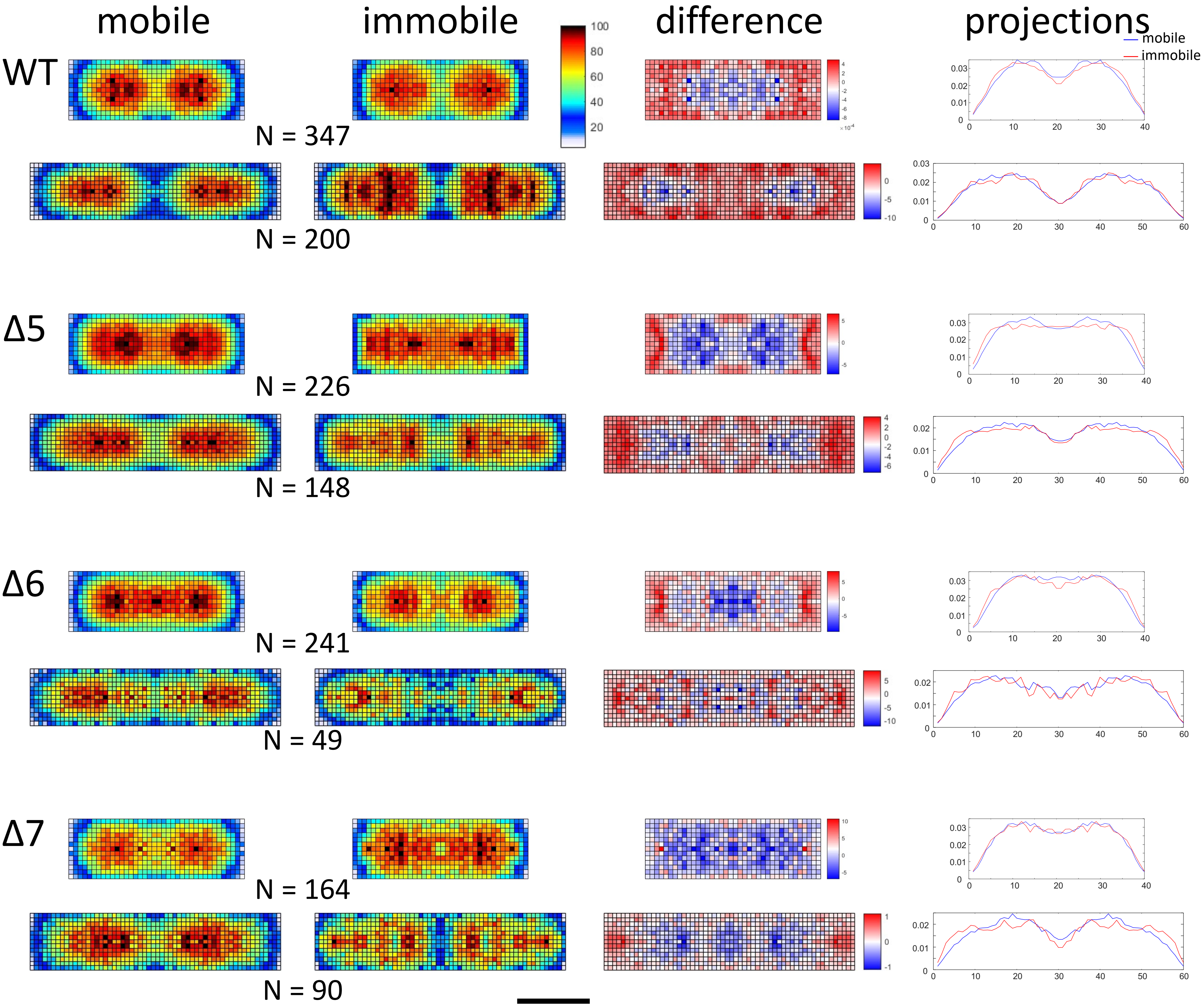


**Fig. S4. Heatmaps of mobile and immobile RNAPs and projections along the long axis for all strains in rich media.** For each strain, cells were sorted into short group (top: 1.8-3.5µm) and long group (bottom: 3.5-4.5µm). Scale bar, 1 µm.

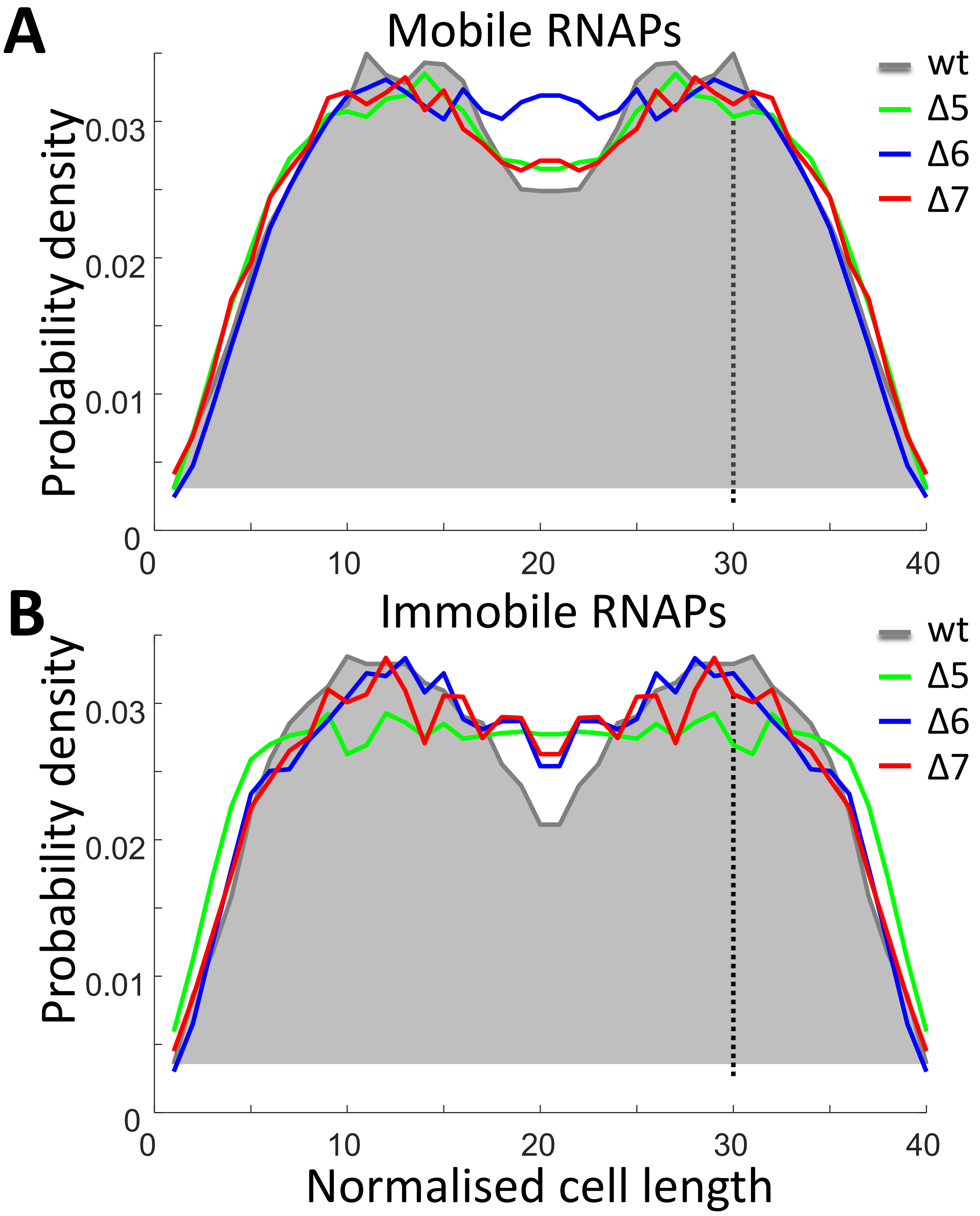


**C**

| Strains | Exterior fractions | |
|---|---|---|
| | Mobile | Immobile |
| wt | 38% | 42% |
| Δ5 | 40% | 44% |
| Δ6 | 37% | 40% |
| Δ7 | 40% | 41% |

**Fig. S5. Spatial distributions of the mobile and immobile RNAPs for short cells in RDM media. A**-**B**. Projections of mobile (**A**) and immobile (**B**) RNAPs along the long axis of the heatmaps for the short-cell group (1.8-3.5 µm length range) in RDM. WT projection is shown as a gray-shaded area. Dashed vertical lines indicate the 25% position along the long axis. **C**. Fractions of RNAPs localised in the exterior 25% region along the long axis.

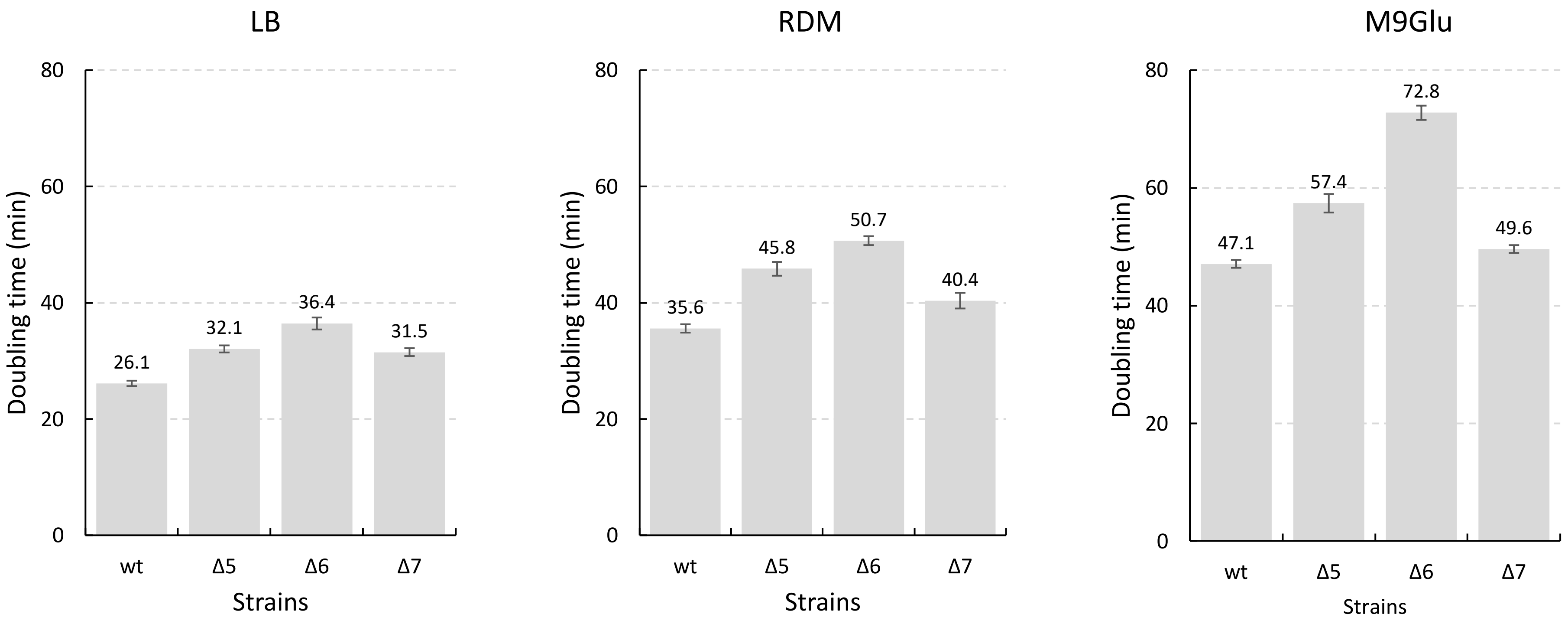


**Fig. S6. Doubling times of wild type and *Δrrn* strains growing at 37°C in different growth media.** Left: LB. Middle: RDM. Right: M9Glu. The means of the doubling times are presented on top of each column; error bars reflect for standard error of the mean (SEM).

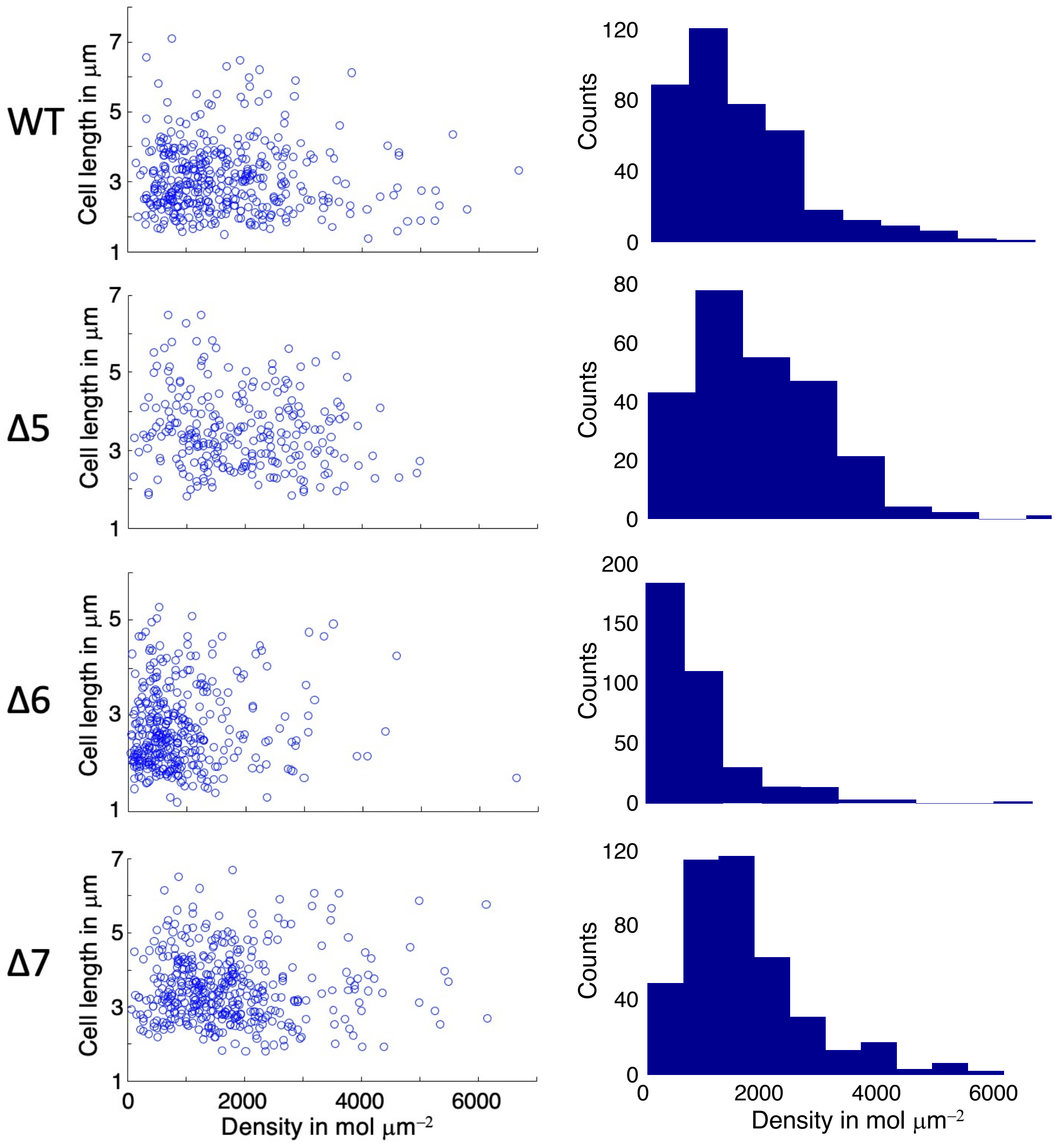


**Fig. S7. Acquired RNAP localizations for all strains in M9Glu media.** Left side: scatter plot of localizations as a function of cell length. Right side: histograms of localisation density.

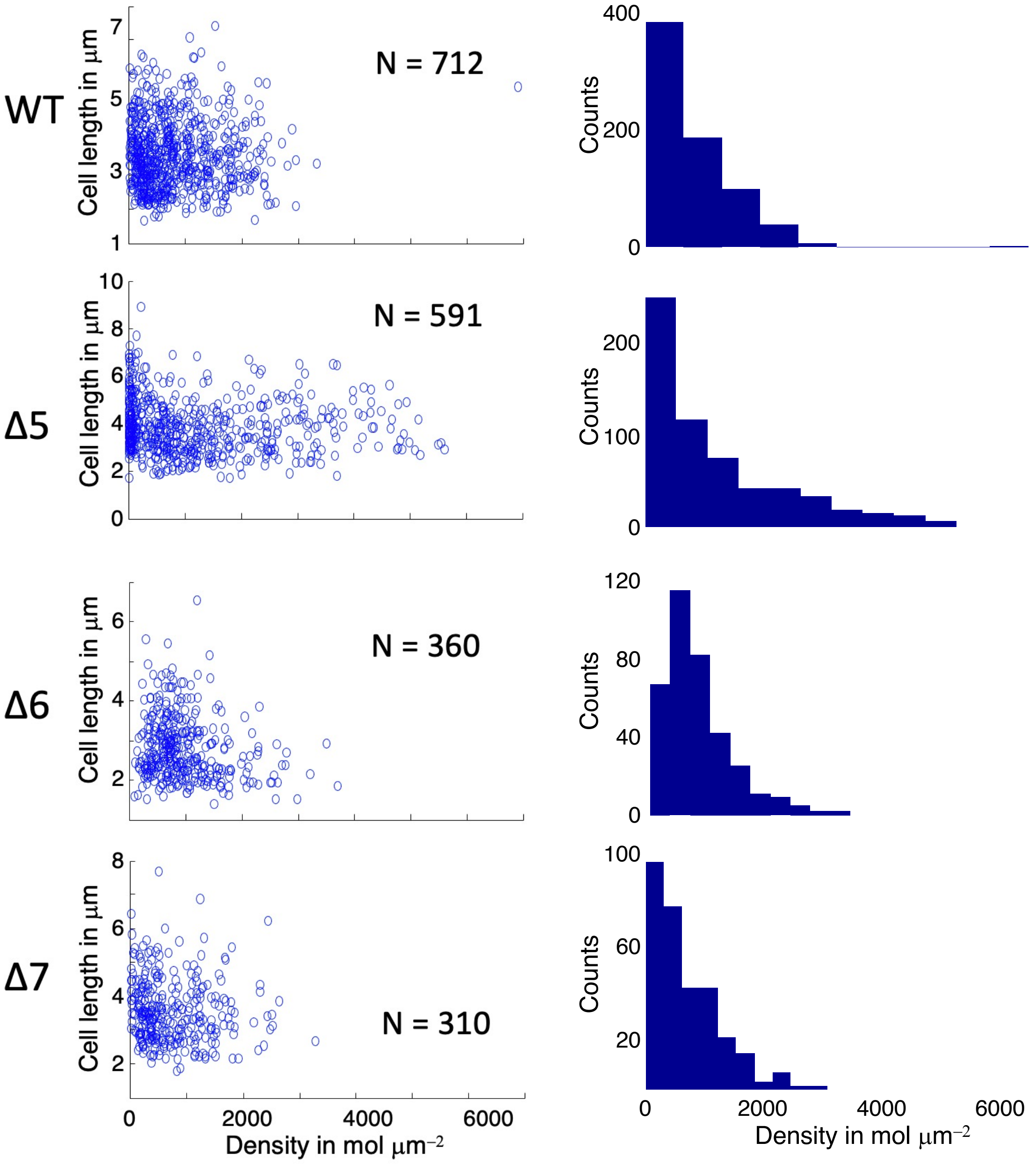


**Fig. S8. Acquired RNAP localizations for all strains in RDM media.** Left side: scatter plot of localizations as a function of cell length. Right side: histograms of localisation density. N denotes number of cells.

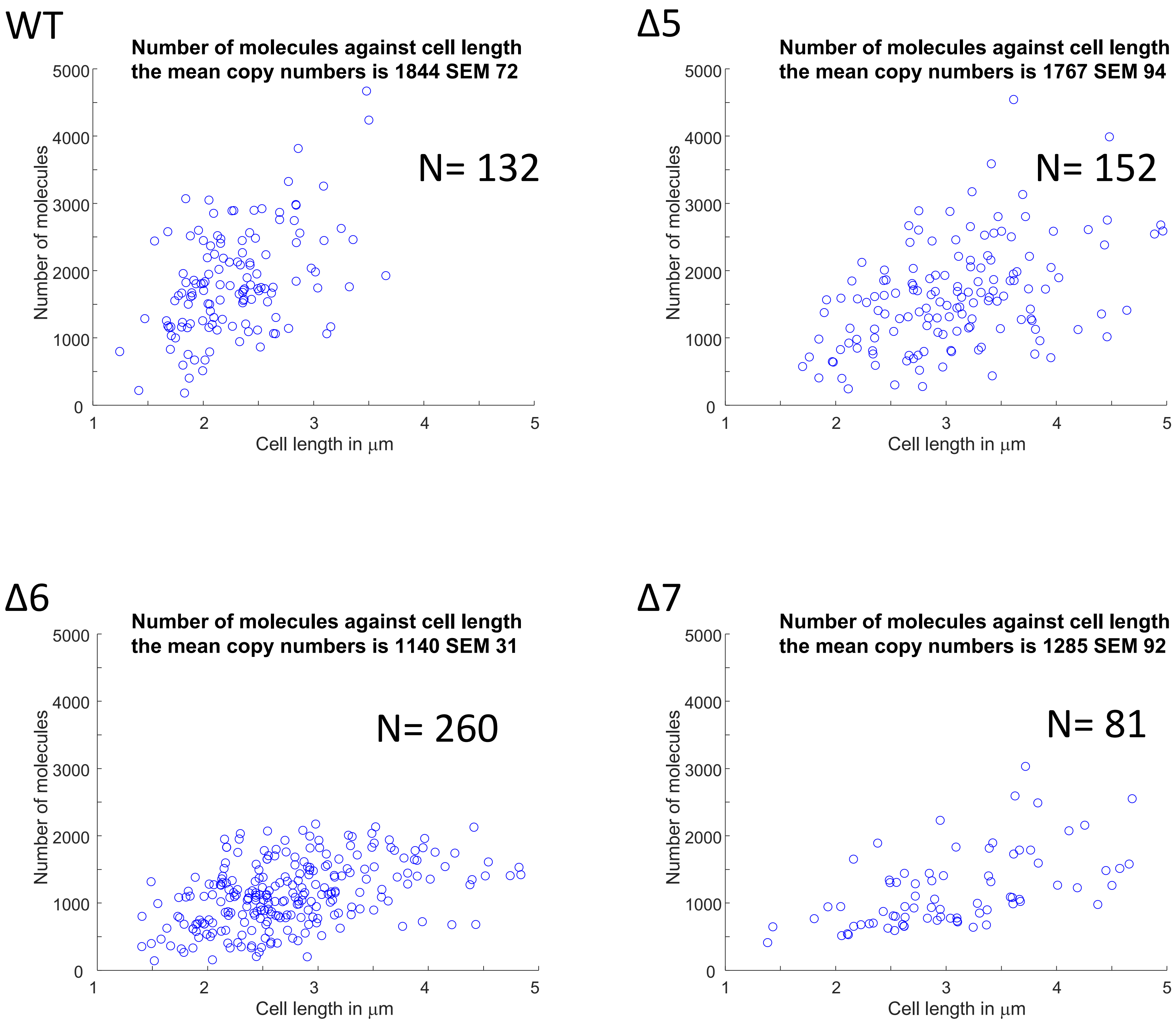


**Fig. S9. RNAP copy number measurement based on PALM measurements in fixed cells in M9Glu media.** The mean copy number and SEM per strain is shown as in the title of each panel. N denotes number of cells.

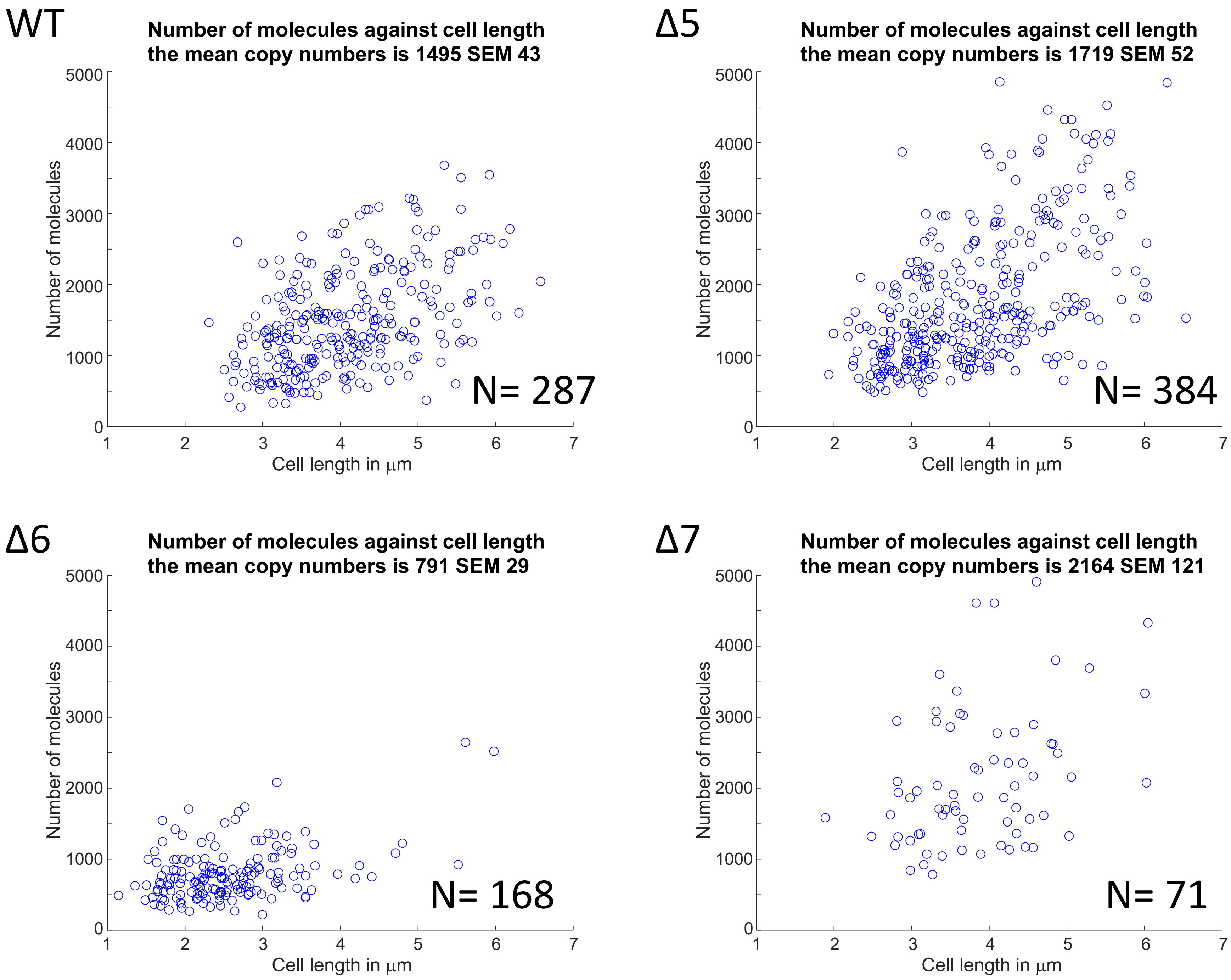


**Fig. S10. RNAP copy number measurement based on PALM measurements in fixed cells in RDM media.** The mean copy number and SEM per strain is shown as in the title of each panel. N denotes number of cells.

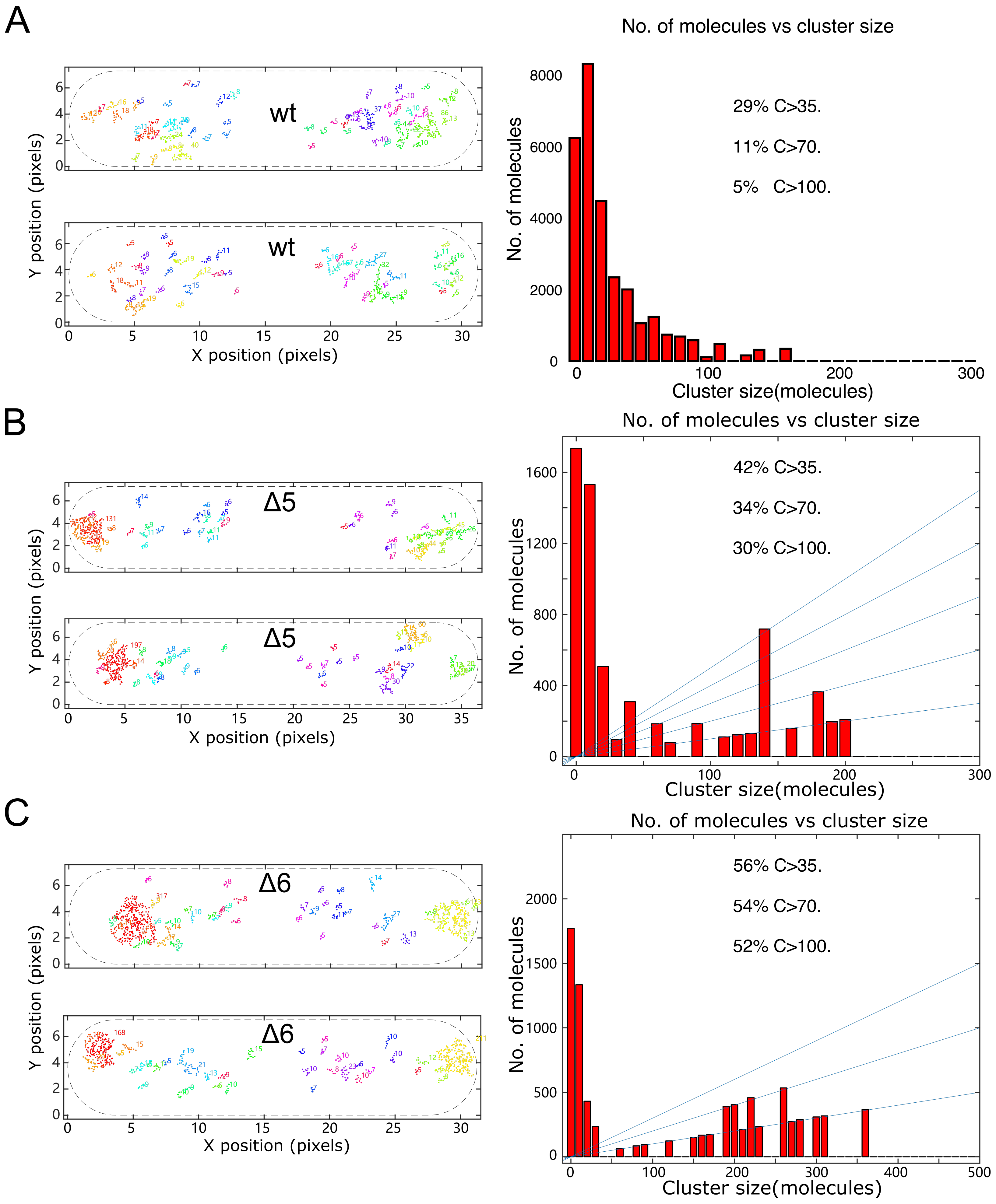


**Fig. S11. Representative cell examples with RNAP clusters by simulation for wt, Δ5 and Δ6 and related clusters histograms.** For a detailed description, see *Methods.* **A-C.** Simulated RNAP localisations (left columns) and frequency histogram for identified RNAP clusters of different size (right columns) in WT cells (panel A), Δ5 (panel B) and Δ6 (panel C).

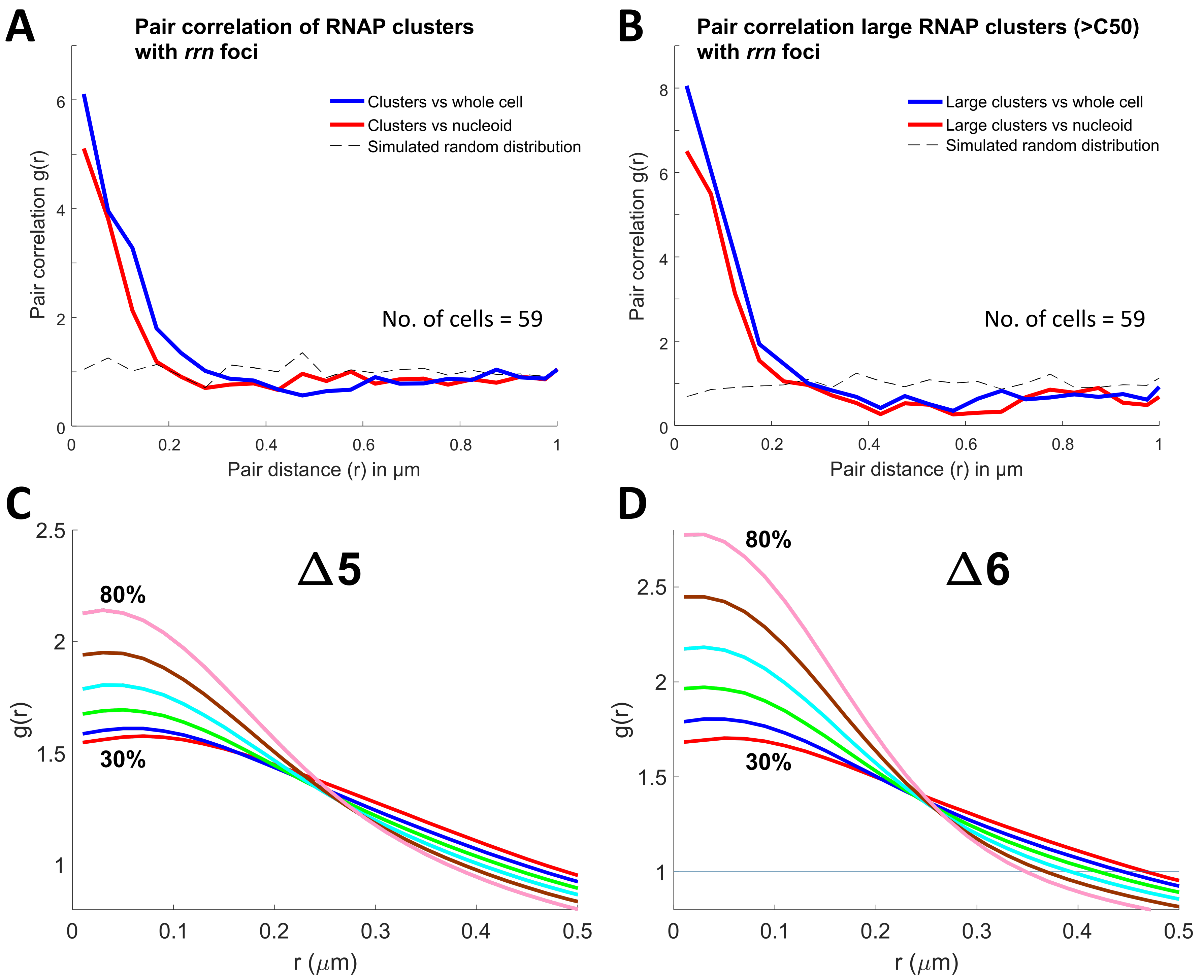


**Fig. S12. Pair correlation of RNAPs clusters with *rrn* foci and simulated data for Δ5 and Δ6.** **A.** Pair correlation of RNAPs clusters in Δ5 in M9Glu media normalised with whole cell (blue curve) and with nucleoid (red curve), compared with simulated localisations throughout the nucleoid. The fraction of clustered RNAPs within 200nm of *rrn* foci: 46.2 ± 2.5 % (SEM) and the fraction of random localisations throughout the nucleoid within 200nm of *rrn* foci 22 ± 1.6 %(SEM). **B.** Pair correlation of large RNAPs clusters (C>50) in Δ5 in M9Glu media normalised with whole cell (blue curve) and with nucleoid (red curve), compared with simulated localisations throughout the nucleoid. The fraction of heavily-clustered RNAPs (>C50) within 200nm of *rrn* foci: 77.3 ± 5.2 % (SEM) and the fraction of random localisations throughout the nucleoid within 200 nm of *rrn* foci 23 ± 1.9 % (SEM). **C-D.** Pair correlation of RNAPs by simulated data for Δ5 and Δ6. Percentages indicate the fractions of immobile RNAPs covered by individual *rrn*. As coverage increases from 30% to 80%, RNAPs pair correlation levels increase.

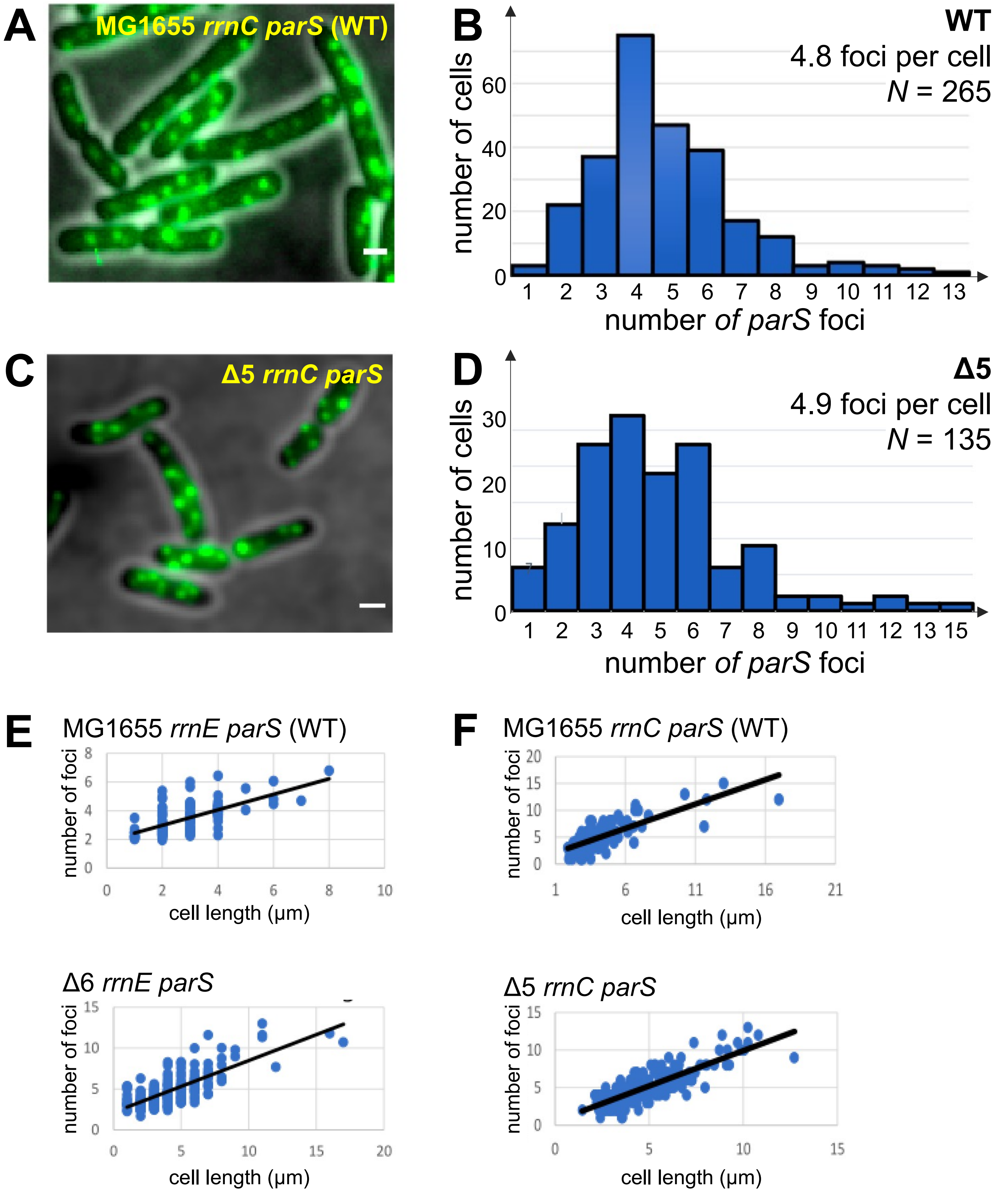


**Fig. S13. Imaging of *rrnC* operons shows increased gene dosage in RDM for Δ5 compared with WT.** Fluorescence imaging of *rrnC* foci in *rpoC:PAmCherry rrnC parS* strains harbouring plasmid pFH2973 expressing parB-yGFP. For details, see *Methods* and Fig. 5. **A.** Example of an image of cells containing parB foci at *rrnC* operons (appearing as bright green foci). Scale bar, 1 μm. **B.** Frequency distribution of the number of foci per cell for WT, along with mean and standard deviation. Cells were segmented and foci were localised and counted using microbeJ (see *Methods* and Fig. 5). **C-D.** Same as in panels A-B, but for the Δ5 strain *rrnC parS*. **E.** Scatter plots showing cell-length dependence of the number of *rrnE* foci per cell for WT and Δ6. Each dot represents a cell; the black line represents the linear fit to the data. **F.** Same as in E, but for the *rrnC*-labelled WT and Δ5 strains.
